# Intrinsic neuronal dynamics predict distinct functional roles during working memory

**DOI:** 10.1101/233171

**Authors:** DF Wasmuht, E Spaak, TJ Buschman, EK Miller, MG Stokes

## Abstract

Working memory (WM) is characterized by the ability to maintain stable representations over time; however, neural activity associated with WM maintenance can be highly dynamic. We explore whether complex population coding dynamics during WM relate to the intrinsic temporal properties of single neurons in lateral prefrontal cortex (lPFC), the frontal eye fields (FEF) and lateral intraparietal cortex (LIP) of two monkeys (*Macaca mulatta*). We found that cells with short timescales carried memory information relatively early during memory encoding in lPFC; whereas long timescale cells played a greater role later during processing, dominating coding in the delay period. We also observed a link between functional connectivity at rest and intrinsic timescale in FEF and LIP. Our results indicate that individual differences in the temporal processing capacity predicts complex neuronal dynamics during WM; ranging from rapid dynamic encoding of stimuli to slower, but stable, maintenance of mnemonic information.

---

Single neuron dynamics in higher cortical areas are heterogeneous, complicating interpretation of their functional roles during cognitive tasks ^1, 2^. A prominent example illustrating this principle is working memory (WM), a mental process strongly associated with the prefrontal cortex (PFC)^3–5^. During WM, information about a transiently encoded stimulus needs to be stored and kept available over a short delay before a response can be made^6, 7^. The neural mechanisms by which this kind of cognitive stability is achieved remain a matter of scientific debate^8–10^.

A large body of experimental studies emphasize a coding scheme whereby information is maintained through persistent firing of single neurons^11–14^. This view is supported by theoretical models demonstrating that persistent activity can emerge from either intrinsic cell properties^15, 16^ or reverberations in recurrently connected populations of selective neurons^17–21^. However, other studies highlight heterogeneous temporal tuning profiles in a majority of recorded neurons^22–24^ or distinct bursts of activity^25^, resulting in a highly dynamic population code underlying WM^26–28^. Supported by theoretical models ^29–31^, those observations have led to the view that maintenance of stable mental representations in WM is possible in the absence of persistent activity in single cells, either through coordinated transient dynamics ^31^ or rapid connectivity changes^30, 73^.

Given the frequent observations of both persistent and dynamic coding at both the population and single neuron level, it is unlikely that these ideas are mutually exclusive^8, 32^. In fact, recent studies have shown that stable population coding can coexist with heterogeneous neuronal dynamics^28, 33^. While those studies stress stable population coding despite overall neuronal dynamics, critically, both regimes interact during categorizations performed by monkeys during WM tasks, relying on both persistent and dynamic neurons^34^. Both transient and persistent activity in single neurons seems to be important for WM. However, the source and function of the neuronal tuning and temporal variability underlying WM population dynamics remain poorly understood.

One of the most striking principles observed across the mammalian cortex is the hierarchical organization of receptive field size. While this phenomenon has been well studied in the visual system, where spatial receptive field size increases with functional hierarchy^35^, a similar principle is currently being uncovered for the temporal domain^36^. Murray et al. (2014)^37^ estimated decay time-constants for neurons’ spiking autocorrelation during baseline ‘resting’ activity, coined ‘intrinsic timescale’, across areas in monkey cortex. They reported that intrinsic timescales increased along the cortical hierarchy. This observation was further supported by evidence from previous studies in monkeys and mice^38, 39^ and results from human imaging studies^40, 41^. Importantly, a recent modelling study demonstrated that the observed gradient of intrinsic timescales arises naturally in a large scale brain model through the balance of intra-and inter-area connection densities^42^.

Intrinsic timescales can be interpreted as the duration over which cells integrate information^43^. According to this view, shorter timescales in sensory areas promote rapid detection of dynamic stimuli^44, 45^, increasing the temporal dimensionality of neural coding. Conversely, longer timescales in prefrontal areas could support integration of information over longer periods of time and could improve signal to noise ratio needed for decision making or WM^43, 46^, but at the cost of dimensionality in the temporal domain. Consequently, heterogeneity in intrinsic timescales found within a cortical area might reflect functional specializations of individual neurons according to temporal processing demands. In line with this, a recent study by Scott et al., (2017)^47^ showed that the fronto-parietal network encodes accumulated evidence with a diversity of neuronal timescales. A similar observation was made by Bernacchia et al. (2013)^48^, who found a reservoir of timescales for reward integration across cortical neurons. Furthermore, orbitofrontal cortex (OFC) neurons associated with a relatively long intrinsic timescale where shown to carry chosen value signals over longer time periods than their short timescale counterparts^49^. Similarly, Nishida et al. (2014)^50^ showed that a more stable baseline activity was associated with higher firing rates during the delay period of a WM in lateral intraparietal cortex (LIP).

Here, we explore how the temporal stability of individual neurons at rest (i.e., pre-trial fixation period) relates to the heterogeneity in population dynamics observed during the subsequent WM trial. We tested whether a neuron’s intrinsic timescale determines its functional role during WM in three brain regions: the lateral PFC (lPFC), the frontal eye fields (FEF) and the LIP. We found that neurons associated with a relatively long intrinsic timescale carried more information about task relevant features than short-timescale neurons. Importantly, in prefrontal areas, long-timescale cells also carried information in a more stable way. This was observed for each task epoch, and was especially striking during the delay period, for both individual cells and at the population level. Additionally, lPFC cells with shorter intrinsic timescales signalled item information earlier during memory encoding and showed richer dynamics during the delay period, suggesting specific functions for distinct populations during individual epochs of a WM task. Lastly, we found that functional connectivity at rest was correlated with intrinsic timescales in FEF and LIP but not in PFC, potentially implicating different mechanisms determining a neuron’s temporal processing characteristics.

## Results

Spikes were recorded from lateral prefrontal cortex (lPFC, n=583), frontal eye fields (FEF, n=323) and lateral intraparietal cortex (LIP, n=281) (Fig. 1a, brain schematic) while monkeys performed a delayed change detection WM task (Fig. 1a, left panel). To initiate a trial monkeys had to fixate a red circle in the middle of a black screen (fixation period: 500ms). Upon successful fixation monkeys were presented with a sample array for 800ms (sample period). A sample array could contain two to five colored squares (items) distributed over six locations (three in each visual hemifield). After a memory delay of 800-1000ms (delay period) a test array appeared on the screen. The test array was equal to the sample array except for one of the items (the target), changed color. Monkeys had to indicate the location of the target with a saccade. For details regarding the task structure refer to Methods and to Buschman et al. (2011)^51^.

**Figure 1:**
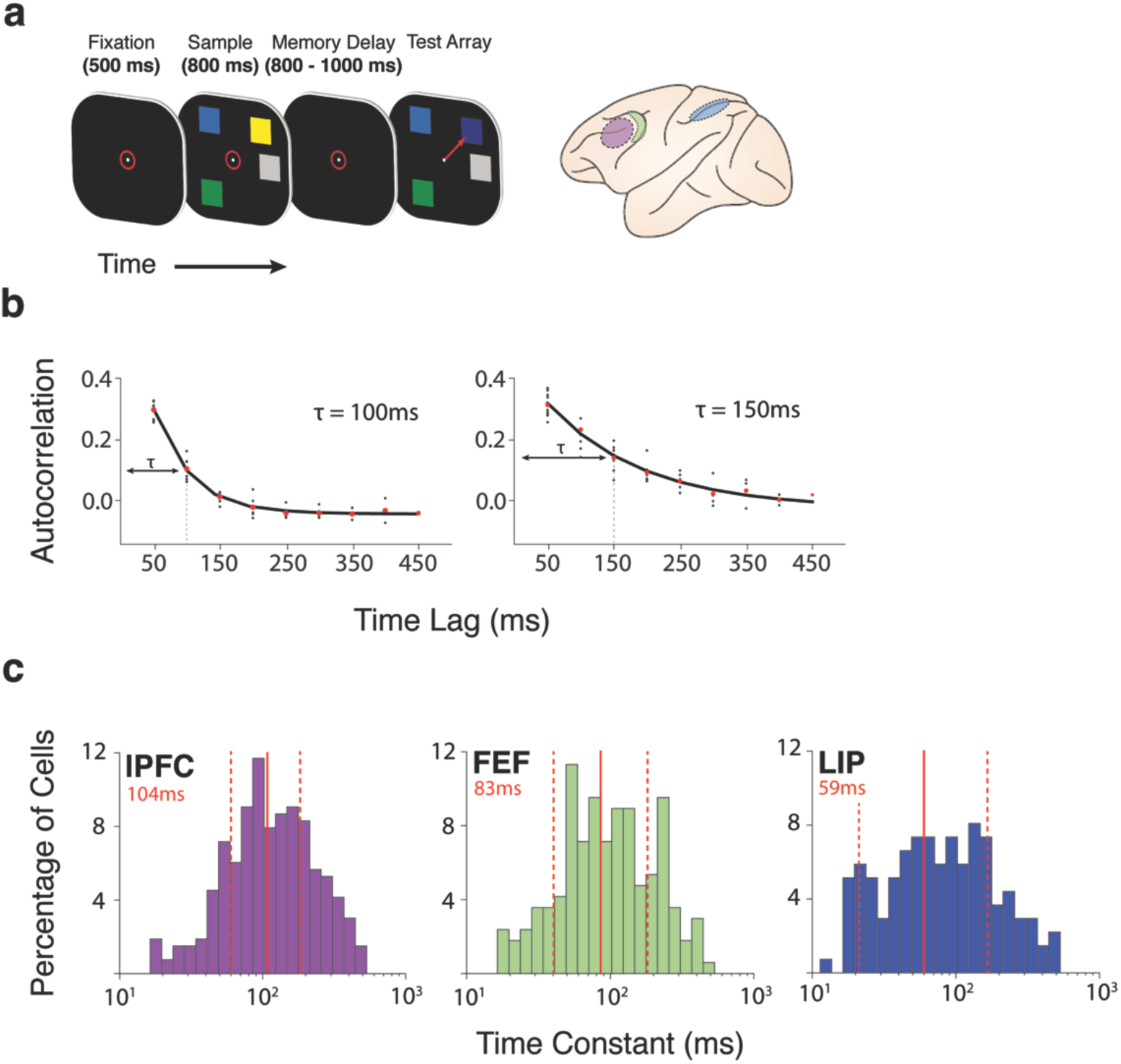
Experimental paradigm & estimation of intrinsic timescales: **a)** Delayed change detection task: Subjects fixated to initiate the start of a trial (red circle, 500ms). A sample array was presented for 800ms consisting of two to five items. After a memory delay (800-1000ms) a test array was displayed that was identical to the sample, except one item (the target) had changed color. Animals indicated target location by a saccade (see ^51^). The left panel shows recording sites: lPFC (purple); FEF (green) and LIP (blue). **b)** Autocorrelation decay functions of two example cells estimated over the duration of the fixation period (left: low τ; right: high τ). The red points denote the mean autocorrelation for a specific time lag. Solid black lines denote exponential fits to the red points. **c)** Histograms of the distribution of intrinsic timescales estimated separately for each cell and each brain region lPFC (n=265); FEF (n=168); LIP (n=136). Color code corresponds to the brain schematic. Red vertical lines denote mean and standard deviations, estimated from log transformed τ values (mean values are displayed in red).

### Defining neural populations by their intrinsic timescale

Our main question was how the intrinsic temporal stability of a neuron^37^ contributes to the dynamic processes underlying a WM task. To investigate this question, we computed the decay time constant of each neuron’s autocorrelation function during the fixation period^37, 42&49^. The resulting quantity is referred to as the intrinsic timescale (tau, τ). A relatively long τ indicates stable firing during fixation, as opposed to a short τ indicating more dynamic baseline firing patterns. Figure 1b shows two example cells from lPFC with different τ’s. We plotted each cell’s τ as a function of brain region (Fig. 1c). Average τ values differed across brain regions (Kruskal-Wallis, p<0.0001). Intrinsic timescales in lPFC were larger than in LIP (Kruskal-Wallis followed by Dunn’s test, p < 0.001), but not FEF (p<0.08). Average τ values in LIP and FEF were not significantly different from each other (p = 0.14). Importantly, we also found that our observed τ values did not depend on the average fixation firing rate in any of the brain regions. (Spearman rank correlation of τ with firing rate; lPFC: r=0.06, p=0.3; FEF: r=0.12, p=0.1; LIP: r=0.08, p=0.3).

### Cells with larger intrinsic timescales carry more WM-related information

To characterize the influence of individual intrinsic firing stability on task involvement, we first quantified the amount of information each cell carried about the task relevant features (i.e., memory item). Here, we used the percentage of variance explained (ωPEV) statistic to measure the extent to which the variability in neural firing rate was determined by color and location^51^.

To determine the relationship between intrinsic timescale and single cell item selectivity, we sorted cells by τ and plotted their respective ωPEV over time (Fig. 2a). Visual inspection of the sorted neural populations reveals a clear pattern towards stronger item encoding as a function of increasing τ in all three brain regions. Splitting the neural population according to their respective median τ value and averaging the ωPEV for each median split draws out this conclusion (Fig. 2b). In lPFC, long τ cells carried more information, especially during the delay period (cluster based permutation test; p=0.003), while in FEF long τ cells were more informative throughout both the sample and delay periods (p=0.008, p=0.02). Interestingly, there was little difference between cell types during the initial transient responses associated with memory encoding, or the response probe at the end of the trial: the functional benefit for long τ cells only emerges during the more stable task epoch. LIP cells showed a similar trend during the sample period, but not during the delay period, where mnemonic coding was poor.

**Figure 2:**
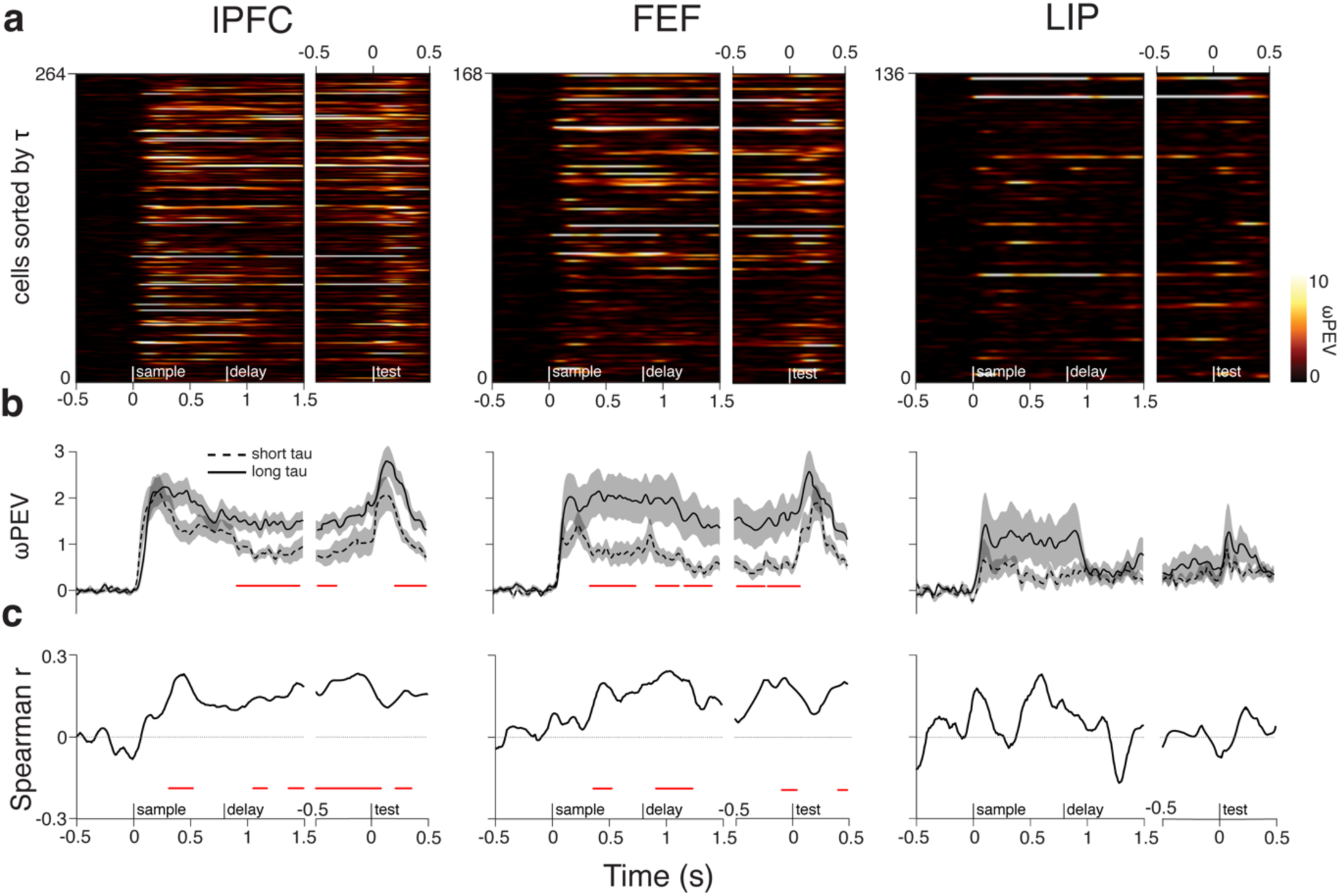
Item information and intrinsic timescales. **a, b & c: a)** Cells sorted by their intrinsic timescale. The color scale indicates the amount of item information i.e. ωPEV present at each time point in a given cell. Traces are smoothed with a Gaussian kernel (s.d. = 50 ms) along the time axis and nearest neighbor interpolation along the cell axis. Initially, trial time is shown with respect to sample onset. However, the duration of the memory delay is variable (800-1000), therefore, we show the last part of the trial (late memory delay until the execution of behavioral response) separately, with trial time shown in reference to presentation of the test screen. The same convention is used for all brain areas (lPFC: n=265; FEF: n=168; LIP: n=136). **b)** Median splits of cells according to their intrinsic timescales. The black dashed line denotes the time resolved average over all cells belonging to the first half of the split i.e. short τ. The black solid line denotes the time resolved average ωPEV over all cells belonging to the second half of the split i.e. long τ. The shaded area represents the standard error of the mean (s.e.m.). The red solid line marks a significant difference between the two splits. **c)** Spearman rank correlation of the intrinsic timescales and the time-resolved ωPEV. The red solid line marks a significant correlation coefficient. Correlation time-courses were smoothed with a Gaussian kernel (s.d. = 50ms); statistics were computed on unsmoothed traces.

To further investigate the observed relationship, we computed a rank correlation between each cell’s τ and ωPEV value for each time point in the trial (Fig. 2c). The positive correlation time courses confirm our results. In lPFC, the correlation between τ and ωPEV shows an initial, transient bump (cluster based permutation test; p<0.01), as also seen in the median split plots (Fig. 2b). This is followed by a progressive increase during the delay period (p=0.0001). In FEF we observe fast rising correlation values during the sample period, which stay high throughout the task (p=0.002). In LIP the correlation between PEV and τ is less straightforward, showing a trend for larger correlation values during the sample period of the task (no cluster survived thresholding), followed by a drop off during the delay period. Although not directly correlated with τ, the average firing rate during fixation might still influence the relationship between τ and the PEV. Therefore, we regressed the average PEV estimated from the periods of significant Pearson correlation (Fig. 2c) against τ factoring out average fixation firing rate (lPFC: β=0.19, p=0.004; FEF: β=0.25, p=0.008; LIP: β=0.2, p=0.017).

### Intrinsic timescale predicts encoding onset during the sample period in lPFC

Cells associated with longer τ values carried more information, which was especially prominent during the delay period in lPFC. In the visual system, smaller receptive fields are useful for detection of rapid fluctuations in stimuli^44^. Analogous, we considered whether short τ cells might therefore be faster at encoding at the time around stimulus onset, since such a division would point towards different functional subpopulations for fast perceptual encoding and longer-term storage of the stimulus.

To investigate this hypothesis, we limited our analysis to cells that showed significant item encoding (ωPEV) at any point during the trial (Supp. Fig. 1). Visual inspection of the average ωPEV in short and long τ subpopulations suggests that low τ cells might encode more information earlier during the sample period in lPFC (Fig. 3a; no cluster survived thresholding). To examine this in more detail, we computed the fraction of currently significant cells for each time point while averaging over τ values for this fraction (Fig. 3b). We found that, especially during the early period of the sample epoch, as more cells became task-selective, the average τ value for those task-engaged cells increased as well. This observation was complemented by a significant positive correlation between τ and encoding onset time (time of first significant ωPEV) during the time period in which the fraction of encoding cells reached 80% percent of its maximum (first 200ms of the sample period) (Spearman rank correlation: r=0.36; p<0.001). This correlation remained significant even after taking the whole sample period into account (r=0.16; p=0.03). This result clearly shows that cells with a shorter τ actively encode task information earlier during stimulus presentation in lPFC. In FEF and LIP we found no such relationship (Supp. Fig 2).

**Figure 3:**
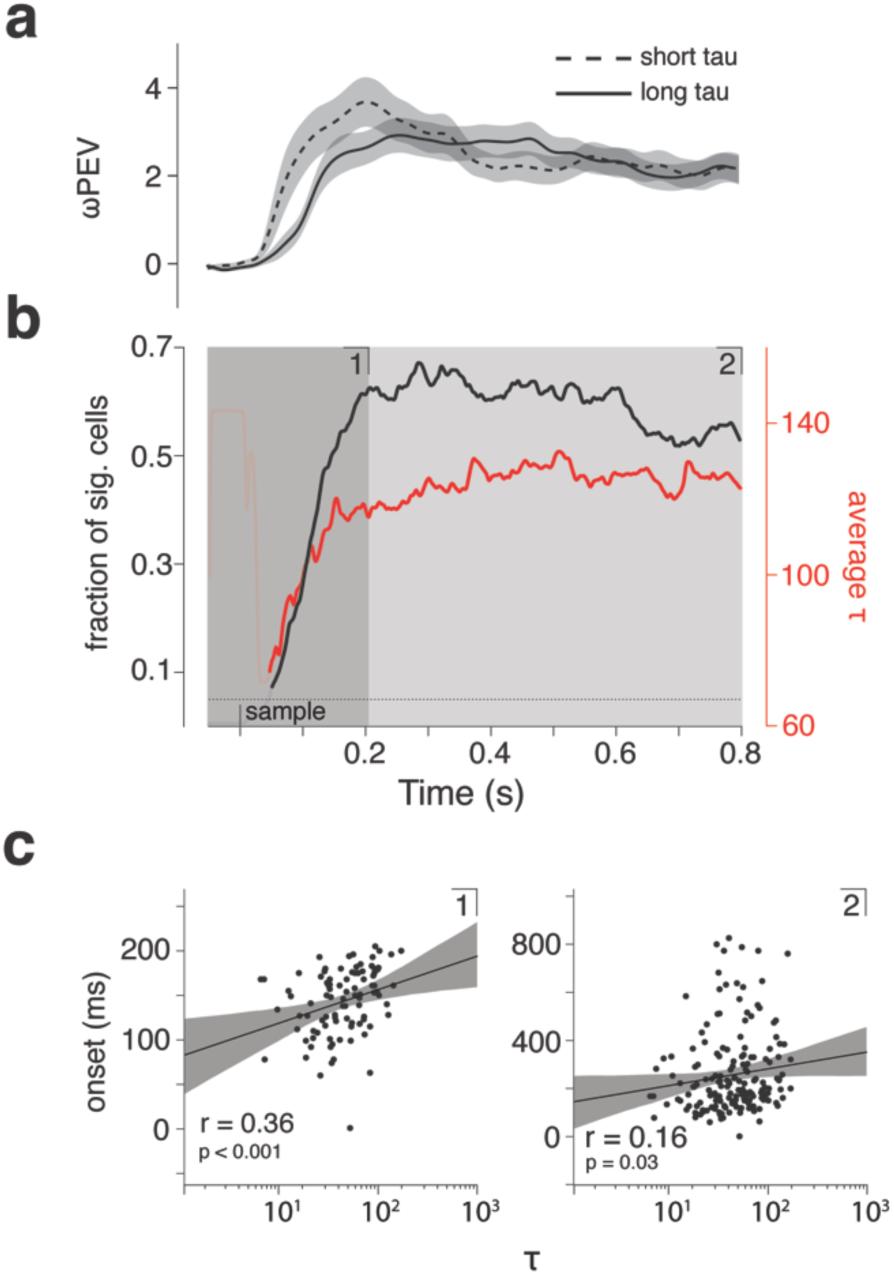
Evaluation of sample period onset times in relation to intrinsic timescales for lPFC. **a**) Average ωPEV per median split (dashed: short τ; solid: long τ) for sample period only; Shaded region indicates s.e.m.. **b)** Black line indicates the fraction of significant cells (ωPEV; p<0.05) at each time point from all cells showing a significant ωPEV at any point in during the sample period. The red line shows the average τ value of cells having a significant ωPEV at that time point. The shaded region marks the time period at which the fraction of significant cells first reached 80% of its maximum value. Occluded parts mark the first time period with fewer than ten significant cells. Numbers in the upper part of the panels refer to **c**. **c**) Left panel: Scatter plots shows onset times (ms) of significant ωPEV estimated during time period **1** i.e. until the fraction of significant cells reached 80% of its maximum value, versus log transformed τ values (Spearman rank correlation: r=0.36; p<0.001; n=96). Each dot marks a cell. Right panel: The same scatter plot but with onset times estimated from the entire sample period **2** (Spearman rank correlation: r=0.16; p=0.03; n=176).

### Population analysis reveals a more stable code for cells with higher intrinsic timescales

Next, we extended our analysis to investigate how τ influences coding dynamics on the level of the neuronal population. Recent studies successfully used cross temporal pattern analysis to characterize dynamics during a range of WM tasks^27, 28&33^. Here, we applied the same method to investigate potential differences in our two subpopulations of neurons (short vs long τ). In brief, we split the observed trials into two independent halves and computed the averaged firing rate, per condition, per neuron, for each half. We then computed all pairwise differences within each split, and correlate the pattern of pairwise condition differences of the whole population at each time point **t** in split one with every time point **t’** in split two, yielding a two-dimensional matrix representing a discriminability score (see Methods). If the discriminating pattern is stationary over time, the within-time correlation values should resemble the between time-point ones; i.e. the pattern should cross-generalize over time. Conversely, a dynamic pattern will result in correlation values lower than the corresponding time-points along the diagonal of the matrix.

To probe whether our previous univariate observations generalize to the population, we first applied the above described decoding approach within-time i.e. for pairs of same time-points between splits (Fig. 4a). Across regions, within-time discriminability time-courses strongly resembled those from the ωPEV analysis (Fig. 4a). In lPFC, short τ cells initially showed a higher discriminability score (cluster based permutation test; p<0.01) which decayed during the delay period, whereas mnemonic information was more strongly represented by long τ cells (p<0.01). In FEF, long τ cells showed higher discriminability throughout the task (p<0.01) while in LIP, item information was only discriminable during the sample period. Here, again, long τ cells showed higher item discriminability than short τ cells (p<0.01).

**Figure 4:**
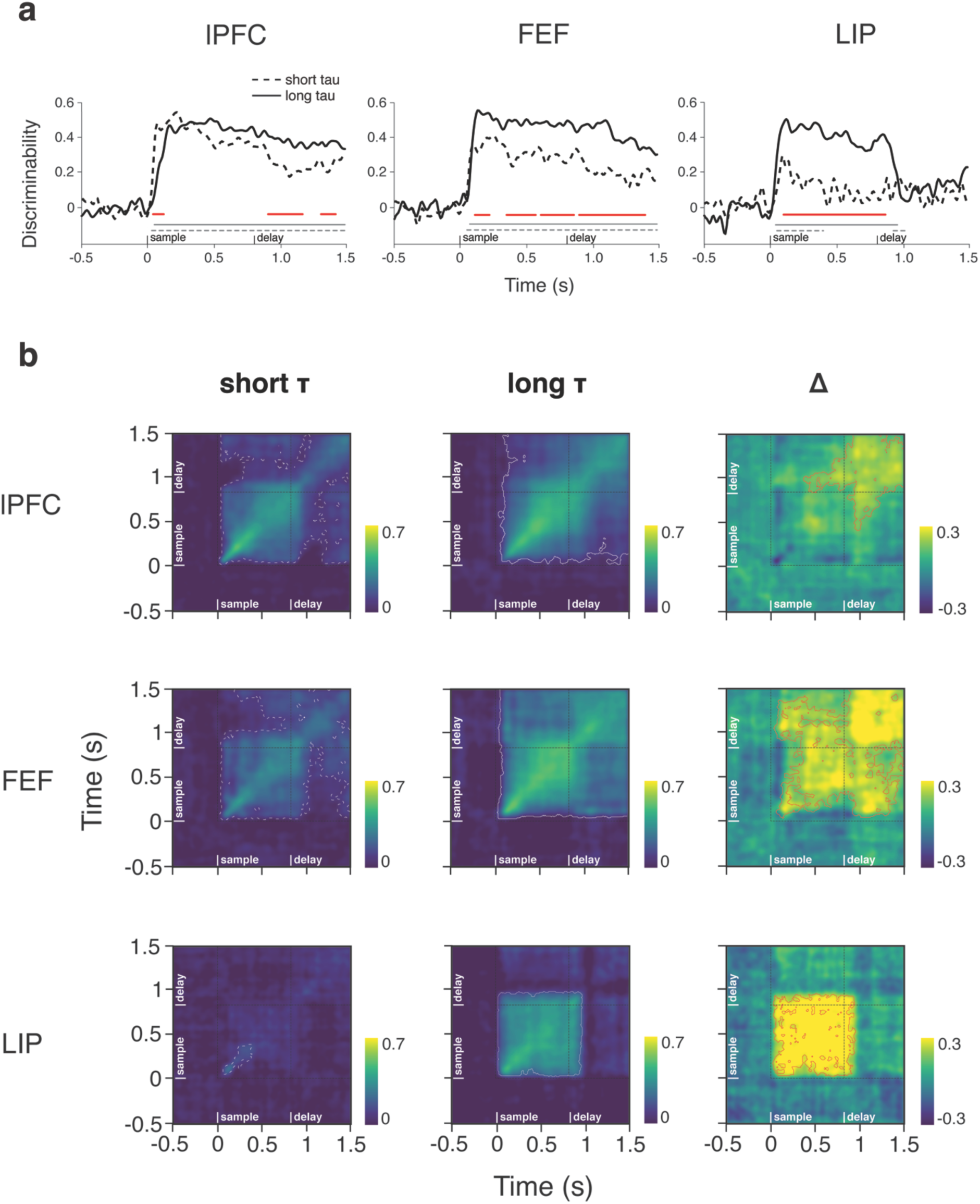
Within-time & cross-temporal decoding and intrinsic timescales. **a)** Within-time decoding. Solid black lines indicate decoding discriminability for long τ cells. Dashed black lines indicate decoding discriminability for short τ cells. Grey lines indicate significant discriminability (cluster based permutation test; p<0.001). Red lines indicate significant differences between the short and long τ cells (cluster based permutation test; p<0.01). **b**) The cross temporal discriminability is color coded. Yellow colors indicate strong decoding while blue colors indicate weak or no decoding. White dashed and white solid contours indicate significant crosstemporal discriminability within respective subpopulation (cluster based permutation test; p<0.001). Fine dashed lines distinguish task periods. First column: short τ subpopulations (dashed). Middle column: long τ (solid). The right column shows the difference between the two first columns i.e. long τ - short τ. Red contours mark a significant difference (cluster based permutation test; p<0.01l) PFC (n=265); FEF (n=168); LIP (n=136).

Figure 4b shows the across-time extension of our decoding approach. Note, the diagonals of the respective matrix plots correspond to time-courses in Fig. 4a. In general, cross-temporal population coding was more prominent and temporally stable for the long τ subpopulation. In lPFC, white contours indicating a significant discriminability score (cluster based permutation test; p<0.001) extend over the entire task duration for the long τ subpopulation (Fig. 4b). In contrast, in the short τ subpopulation off-diagonal discriminability drops to chance level at the transition between sample and delay periods and also within the delay period. Subtracting the discriminability matrices computed for the two subpopulations (long τ - short τ) reveals significantly stronger and more stable decoding during the delay period within long τ cells (cluster based permutation test; p<0.01). In FEF, the discriminative pattern of the long τ cells strongly cross-generalizes both within task epoch as well as across task epochs (white contours, p<0.001). This strong cross-generalization is absent in the short τ subpopulation which shows significantly less cross-temporal decoding (p<0.01). In LIP differences between τ populations are confined to the sample period where long τ cells show a stronger discrimination score (p<0.01), which is remarkably stable (p<0.001).

To illustrate population decoding for both item features separately, we conducted within and across-time temporal discriminability analysis for location and color only (see Methods) (Supp. Fig. 3 & 4; for statistical test values refer to figure legends). For locations only, differences between decoding in τ subpopulations in lPFC and FEF were similar to those observed for combined item decoding (i.e. color & location; Fig. 4) (Supp. Fig. 3). In LIP, location information was weak but present during the delay period (p<0.05), as evident from the crosstemporal plots. Interestingly, long τ cells showed a similar re-emergent pattern during sample and late delay periods. Apart from a very weak effect in FEF, color information was only present in lPFC (Supp. Fig. 4). Here, the long τ subpopulation showed stronger within and across time discriminability scores, especially during the late sample and early delay periods. Color encoding was only weak in FEF and absent in LIP, preventing an informative comparison between τ subpopulations.

### Intrinsic timescales predict single cell temporal coding dimensionality

So far, our results show the existence of a link between an individual cell’s intrinsic firing stability and the magnitude of its item encoding. However, the PEV does not make any assumption about the consistency of neural responses over time nor does it take into account a neuron changing its selectivity^24, 28&52^. While we show that on the population level, intrinsic timescale affects the stability of information across time, we have no single cell measure quantifying temporal coding stability for an individual cell. Hence, we chose to evaluate the effective temporal coding dimensionality (N_*eff*_; where N stands for the number of principal components) of each cell via principal component analysis (PCA; see Methods). In neural population analyses, PCA is classically used for dimensionality reduction; i.e. each neuron counts as a dimension and each trial or time point counts as a point in neuronal state space. Here, we apply the same method to a single neuron with adjacent time bins counting as dimensions and mean condition differences firing rates counting as points in ‘time state space’. Specifically, a high N_*eff*_ indicates a high temporal dimensionality which can be caused by either non-specific activity, unstable firing or switching selectivity.

We estimated the N_*eff*_ in a stepping 500ms time window composed of 10 independent 50ms bins (step size of 50ms). This particular relation between time window and bin size was chosen since it allowed us to evaluate single epochs of the task while capturing most of the temporal coding dimensionality within a given task epoch (Supp. Fig. 5). As expected, N_*eff*_ was highest during the fixation period, when condition specific firing was absent (Fig. 5a). Eliminating condition specific firing by shuffling condition labels yielded N_*eff*_ similar to those of the fixation period (data not shown). In lPFC and FEF, cells with longer τ values displayed a lower temporal dimensionality during sample and delay periods (cluster based permutation test; p<0.001) (Fig. 5a). In LIP, the difference between N_*eff*_ of the two split halves was significant only around the sample period. The correlation time-courses (Spearman r) in Fig. 5b confirm this observation. In FEF, the correlation between N_*eff*_ and τ steadily increases over time, peaking during the delay period (cluster based permutation test; p<0.01). In lPFC a strong correlation is evident already early during the sample period (p<0.01), which dips between the two task periods, suggesting a τ independent state transition, in order to re-emerge during the delay period (p<0.001). In LIP a dependence of N_*eff*_ on τ is clearly only present during the sample period (p<0.01). Scatter plots of all neurons/area in Fig. 5c show the same correlation but for the average N_*eff*_ estimated from independent time periods within task epochs. Since N_*eff*_ gives an estimate of the temporal variability of neuronal firing, we should expect to observe a higher N_*eff*_ for longer τ values during the fixation period. In fact, we found such a relationship in lPFC (Fig. 5c; Spearman rank correlation: r=-0.15, p=0.013) but failed to observe a significant effect in FEF (r=-0.13, p=0.1) and LIP (r=-0.1, p=0.23).

**Figure 5:**
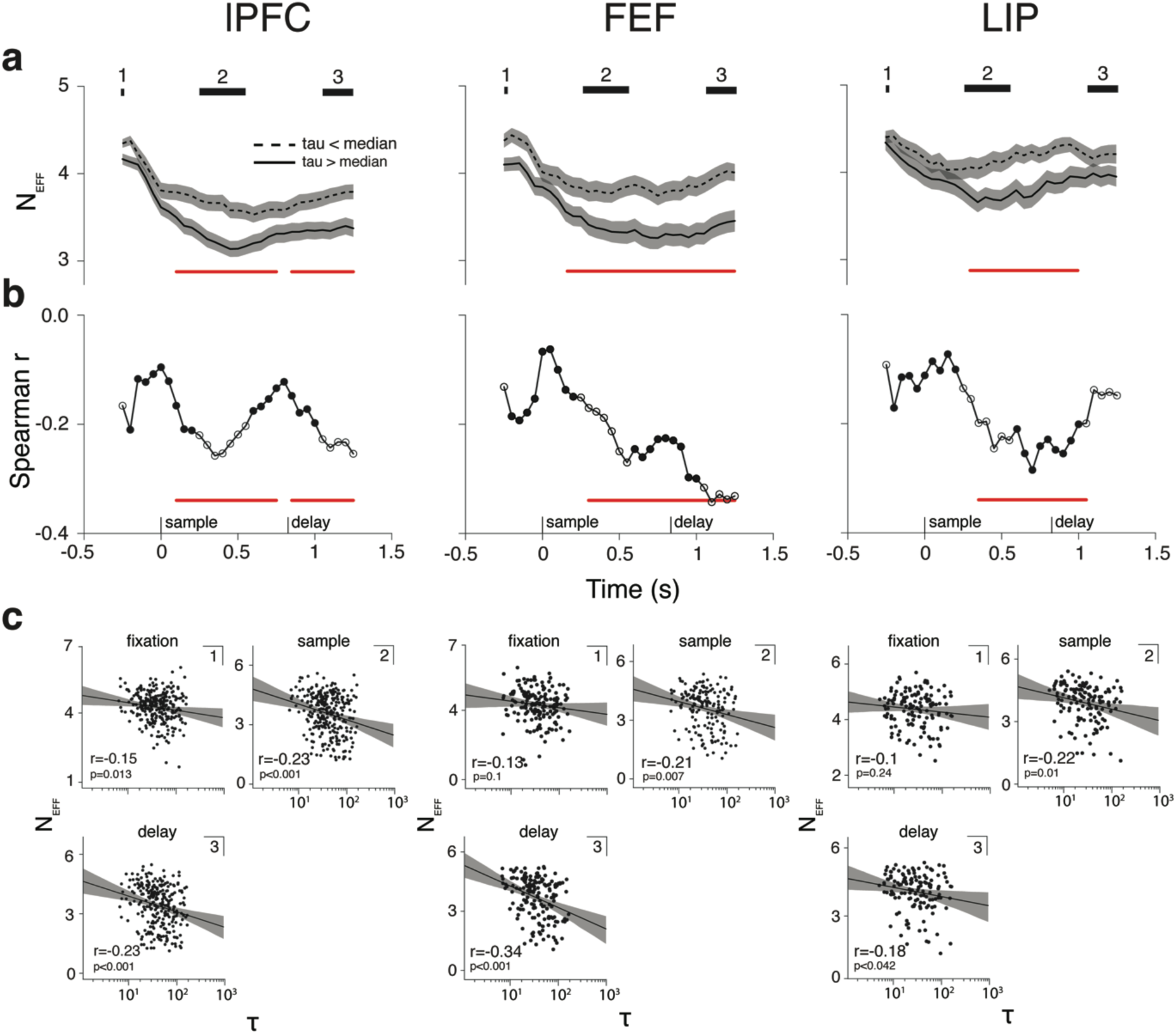
Effective temporal dimensionality and intrinsic timescales. **a)** Effective temporal dimensionality (N_*eff*_) estimated at a single cell level (N_*eff*_ was estimated from a sliding 50ms*10 (500ms) time window shifted by 50ms on each step. Solid black line: Average N_*eff*_ of the respective populations over the course of the trial (left column: lPFC, n=264; middle column: FEF, n=167; right column: LIP, n=134). Dashed red line: Average N_*eff*_ for short cells τ cells. Solid red line: Average N_*eff*_ for long τ cells. Red horizontal line: significant difference between the two splits halves (cluster based permutation test; p<0.01). Shaded regions mark s.e.m.. Black bars mark N_*eff*_ estimated within independent trial epochs (**1**: fixation; **2:** sample; **3**: delay). **b**) Spearman correlation between N_*eff*_ and log transformed τ values. Each dot marks the correlation coefficient as a function of time i.e. the center of the time window from which N_*eff*_ was estimated. Red line: significant Spearman correlation coefficient (cluster based permutation test; p<0.01). Hollow dots mark independent trial epochs (**1**: fixation; **2:** sample; **3**: delay). **C**: Scatter plots for N_*eff*_ estimated from independent trial epochs (number in left corner refers to number on black bar) and τ values. lPFC (fixation: r=-0.15, p=0.013; sample: r=-0.23, p<0.001; delay: r=-0.24, p<0.001). FEF (fixation: r=- 0.13, p=0.1; sample: r=-0.21, p=0.007; delay: r=-0.34, p<0.0001). LIP (fixation: r=-0.1, p=0.23; sample: r=-0.22, p=0.01; delay: r=-0.18, p<0.042).

### Robust temporal dynamics serve as a feature for discrimination

Next, we hypothesized that if a low τ cell genuinely encodes item information, albeit in a temporally dynamic way, the temporal activity pattern should carry information not evident in a purely stable cell. In brief, per neuron, we split trials into two independent halves and quantified the correlation between the temporal pattern of item specific activity estimated over 200ms between the two split halves (see Methods). If the temporal activity trace is corrupted by noise, correlation between the split halves (i.e. the discriminability score) should be small. Similarly, a condition specific but temporally stable trace would also translate into a comparatively small discrimination score. However, a dynamic temporal activity pattern should result in a relatively high discrimination score.

Indeed, item information was discriminable in the temporal coding dynamics inherent in both short and long τ cells (cluster based permutation test, p<0.001) (Fig. 6, for lPFC & Supp. Fig. 6, for FEF & LIP). In lPFC, temporal discriminability time courses were strikingly similar during both the sample and test periods. During the delay period, temporal discriminability initially rose but then stayed significant only for short τ cells (p<0.001), which showed significantly higher decoding during part of the delay (p=0.017). In FEF and LIP we did not observe any significant difference between the two subpopulations separated by τ (Supp. Fig. 6). Interestingly, in all three brain regions, our method seems to capture transitions between epochs as decoding falls back to baseline as task epochs unfold.

**Figure 6:**
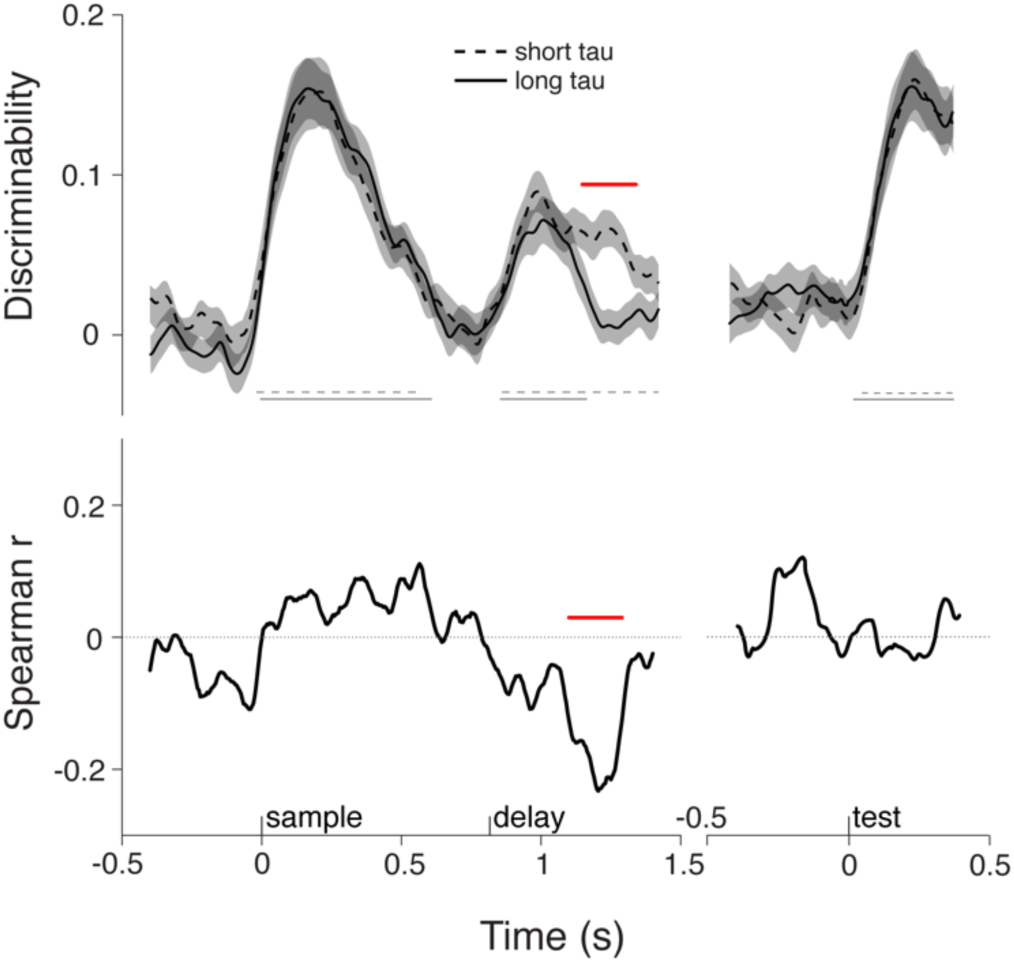
Temporal discriminability in lPFC. The upper panel shows the temporal discriminability score (see methods) averaged over neurons. Black solid line indicate the average over long τ cells and black dashed lines indicate the average over short τ cells. Shaded area indicate s.e.m.. Thin grey lines indicate significant discriminability (cluster based permutation test; p<0.001), for respective τ splits. The red solid line depicts a significant difference between the two splits of the data (cluster based permutation test; p<0.05). The lower panel shows the Spearman rank correlation between τ and discriminability score for each time point. The Solid red line denotes a significant correlation (cluster based permutation test; p<0.05)

Correlating τ and temporal discriminability over time accentuates the observation made from the median splits. In lPFC the correlation time-course shows a significant dip during the delay period (p<0.001) emphasizing stronger temporal discriminability for short τ cells (Fig. 6). Interestingly, a similar dip is evident in the FEF correlation time course at a later time point in the delay (p<0.01) which was only hinted at in the median split plots (Supp. Fig. 6a).

Note, it is improbable for any cell to encode information in a perfectly stable way and strong item encoding is likely to drive much of the observed correlation. However, keeping in mind the weaker item encoding for short τ cells as observed by the ωPEV, an equal or even stronger temporal discriminability score for the same cells strongly speaks for a robust but dynamic coding within this subpopulation.

### Functional connectivity at rest correlates with intrinsic timescales

Next, we tested possible sources leading to our observed τ values. A recent modelling study proposed that intrinsic timescales could, in part, arise through local connection densities^42^. This observation raises the possibility that neurons associated with a long τ value might be part of a local network hub with a high concentration of incoming and outgoing connections^34, 39^. To test this hypothesis, we probed functional connectivity using spike count correlations (rSC), also referred to as noise correlations^54^ (see Methods). Spike count correlations were estimated from the fixation period since we wanted to determine whether τ depended on functional connectivity at rest i.e. before a stimulus synchronized network activity. We found a significant correlation between τ values and rSC when pooling over all brain regions (Spearman rank correlation; r=0.18, p=0.004) (Fig. 7 upper left panel). Spike count correlation can depend on basic firing rate^55^. However, we found no evidence for a significant correlation between rSC or τ and fixation activity, respectively (Supp. Fig. 7; r=-0.05, p=0.37; r=-0.05 p=0.38). Furthermore, the relationship between rSC and τ was also present when controlling for baseline firing using a multiple linear regression model (β=0.03; p=0.007). Together, this implies that differences in fixation activity were not responsible for the positive relationship between rSC and τ. Separating out individual brain regions revealed a more diverse picture: In lPFC, rSC and τ were not correlated (r=0.04, p=0.63, Fig. 7 lower left panel), while both FEF and LIP showed a positive correlation between rSC and τ (FEF: r=0.28, p=0.015; LIP: r=0.33, p=0.013), which held true when controlling for baseline firing using the multiple regression model (FEF: β=0.05, p=0.02; LIP: β=0.04, p=0.02).

**Figure 7:**
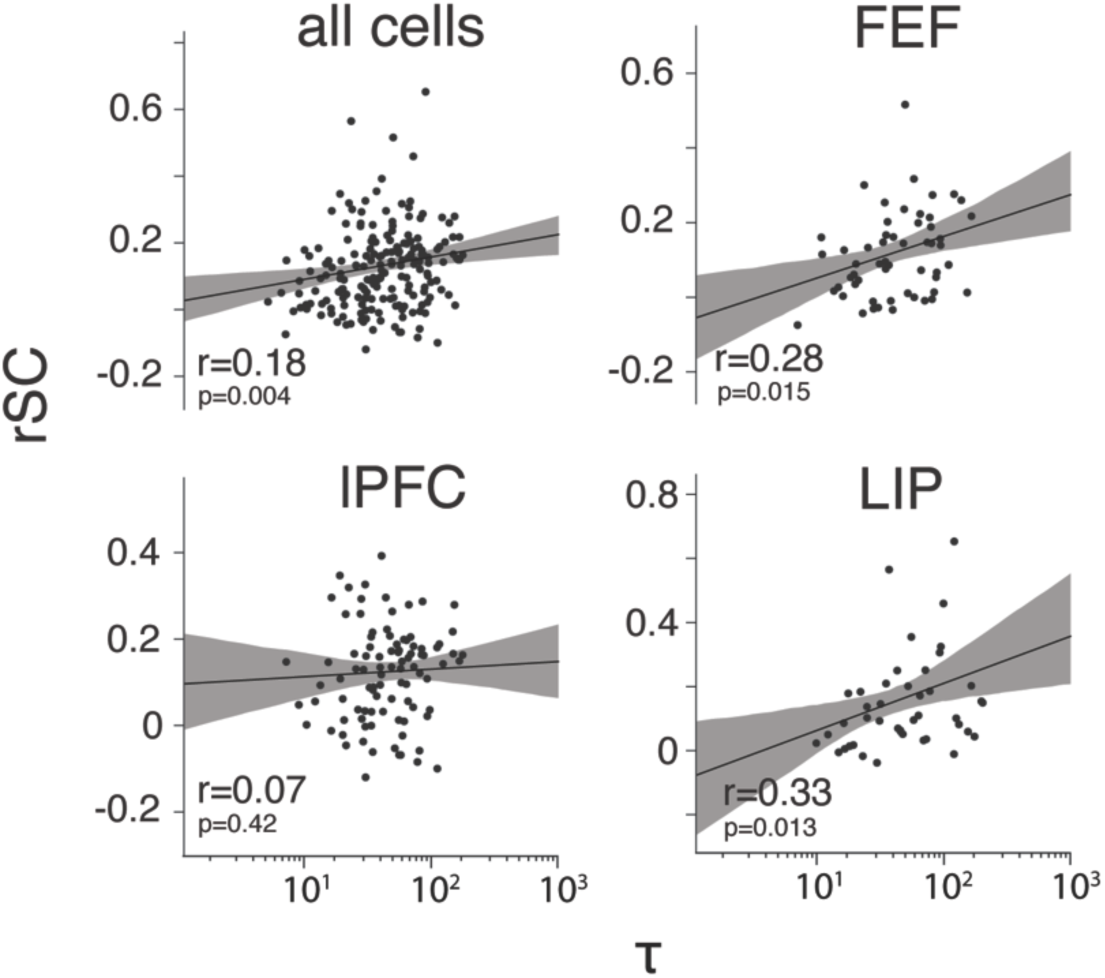
Spike count correlations and intrinsic timescales. Scatter plots for rSC on the y-axis versus the τ values on the x-axis. rSC is estimated during the fixation period and between cells on the same contact. Each dot depicts one cell. The r value within panels depicts the Spearman correlation coefficient and associated p-value, for the two variables. All cells: n=255; lPFC: n=126; FEF: n=74; LIP: n=55. Black solid lines show linear regression lines. Shaded region depicts 95% confidence interval.

## Discussion

Heterogeneity in temporal tuning observed during WM is related to baseline firing dynamics of individual neurons. Specifically, we found that cells with a longer intrinsic timescale carried more information throughout the task epochs, and particularly during the delay period in prefrontal areas. Important to WM function, mnemonic coding of long timescale cells was more stable across time in lPFC and FEF. In contrast, short timescale cells responded more rapidly during encoding in lPFC, and carried rich temporal information. Overall, we demonstrate an intrinsic distinction between functional populations which perform complementary computations: rapid encoding or longer-term storage of information.

The core finding of this study is that the baseline temporal stability of individual neurons in lPFC predicts how robustly they code WM-related information, especially during the mnemonic delay period. Early studies of WM focused on persistent activity in lPFC representing specific items in memory^4^, however more recent studies also reveal more complex dynamics during encoding and maintenance^25, 26&27^. In particular, it has been shown that single cells become active at different time points during the trial, encode information only transiently, possibly multiple times and/or switch their selectivity over time^23, 28&56^. Importantly, we now find that the metric of intrinsic timescales as estimated at rest (i.e., pretrial fixation period) predicts the coding dynamics during WM.

Recent evidence shows that despite transient dynamics during sample and early delay periods, information in the later delay period was maintained in a relatively stable population code^28^. Additionally, a study by Murray et al. (2017)^33^ showed that during WM, stable and dynamic population codes coexist. Conceptually, in those studies, dynamic and stable mnemonic representations refer to orthogonal subspaces in high dimensional neural state space^33^. Here, we extend those observations by proposing that they are linked to intrinsic properties of neuronal subpopulations: long timescale cells more strongly contribute to the stable subspace whereas short timescale cells contribute to the dynamic subspace. Our data also shows that this is mostly the case during the delay period in prefrontal areas. This contrasts with LIP, where encoding did not significantly differentiate between timescale subpopulations during the delay, where mnemonic coding was absent. Rather, larger timescale cells in LIP stably encoded more information during the sample period. A previous study recording from monkey LIP during a simple delayed response task, showed that baseline firing stability modulated stimulus specific activity during the delay period^50^. In that study baseline firing stability was estimated by a relatively coarse metric, namely, the correlation value between spike counts within differentially spaced time bins. This is qualitatively different than measuring the decay time constant of those correlation values over all given time bins. Furthermore, the complexity inherent in our task (color, locations & load effects) as well as relatively poor encoding within the LIP population might have diluted an otherwise clearer relationship. Using location information as discriminative feature revealed some cross-temporal generalization extending to the delay period in LIP (Supp. Fig. 4).

Results from an information theoretic approach suggest WM information can be stored more faithfully if the brain first encodes information appropriately before passing it to persistent activity networks^57^. Within the same study, the authors showed that human performance is better fit by such a two-step model than by a model mimicking direct storage in persistent channels^57^. Here we show that the measure of intrinsic timescale can help explain differences in temporal dynamics and therefore distinct computational roles within a complex WM task. In lPFC we saw that stable coding within the long timescale subpopulation generalizes only weakly across the different epochs of the task (i.e., sample to delay period), suggesting that in lPFC, the neural code transitions between different epochs, ultimately to be stored in a more stable format during the delay epoch through cells with longer intrinsic timescales. This seems to be in accordance with previous observations in PFC^27, 28^ and with the two step model described by Koyluoglu et al. (2017)^57^, while emphasizing distinct networks for perception and storage which are potentially both present during the sample period^52, 58–60^. A similar conclusion had been reached by Murray et al., (2017)^33^, who suggested orthogonal mnemonic and perceptual representations during the sample period with the latter decaying during the delay period. Contributing to the evidence of distinct roles for cells with different timescales, we found that more dynamic cells, as defined by intrinsic timescale, coded for item information relatively early during the sample period. This onset cascade early in the sample period implies a more specific role for short timescale cells since they more rapidly detected the presented stimuli. This result might further play into a two-step model (i.e. rapid perceptual encoding and transfer to a robust stable code) of WM encoding leveraging the presence of a heterogeneous pool of available intrinsic timescales. It is important to note that while discussing different neurons being recruited for certain tasks, those neurons i.e. their respective timescales lie in a continuum rather than belonging to categorically distinct subpopulations. It therefore seems likely that contributions of individual neurons are weighted according to evolving task demands^2^. Along those lines, functional subpopulations are best thought of as distinct dimensions of the neuronal population rather than specific classes of cell types with distinct roles. In FEF, the difference between short versus long timescale cells was striking in terms of magnitude of information encoding during sample and delay period. Different to lPFC, the discriminative pattern within the long timescale subpopulation cross-generalized more strongly from the sample, all the way through the delay period. Note, in our task, strong, temporally stable signals as observed in FEF are unlikely to reflect oculomotor preparation since monkeys can not prepare a response in advance.

High temporal autocorrelation implies temporal stability, which seems well suited for WM. On the other hand, temporal autocorrelation also translates to a lower temporal dimensionality, which necessarily limits the informational capacity of coding over time. A recent, influential study found that high neural dimensionality is crucial for complex behaviour^1^. Specifically, the dimensionality of the neural representation is expanded through neurons exhibiting mixed selectivity in response to task factors, maximising the possibility space for linear classification i.e. readout of the population activity^11^. This view can be extended to the temporal domain, where dynamic changes in neural activity over time i.e. switching selectivity are reflected in a high temporal dimensionality. Indeed, the information potential of a highly dynamic network presentation is directly proportional to the statistical independence between time points^61^. Here, we could show that a faster decay of temporal autocorrelation at rest directly translates to a higher coding dimensionality during the task. While our measure of temporal coding dimensionality (N_*eff*_) captures selectivity to task parameters, it also reflects the temporal stability or instability of the tuning profile, extending the ωPEV measure substantially. This is evident in the clear epoch transitions in lPFC: long timescales cells maintain information in a lower dimensional temporal state only within each task epoch, not in transitions between epoch. Conversely, short timescale cells exhibit a higher temporal coding dimensionality even before the start of the delay period. In FEF the relationship between intrinsic timescale and temporal coding dimensionality steadily increases throughout the task, peaking during the delay period, an observation which again adds on the observed ωPEV time courses. Complementing those observations, we found that robust but temporally dynamic coding can discriminate information which would otherwise be lost in a purely stable encoding. Additional information inherent in temporal dynamics was particularly evident in short timescale cells in lPFC during the delay. Here, short timescale cells showed weaker item encoding as quantified by the ωPEV. Nevertheless, their inherently high dimensional coding properties could be used to decode memory-related information.

Temporal dynamics are especially important within the context of cognitive flexibility. According to the principle of adaptive coding, PFC represents information in a dynamic and therefore flexible manner, where neural resources can be recruited ‘on the fly’, according to behavioural demands^62^. A recent study investigating neural correlates of cognitive flexibility in monkeys found that neurons effectively switching their selectivity over time were most active when cognitive control was demanded^63^. Crucially, most of the variance at the population level could be attributed to time or the interaction between time and trial type, showing that different cue-probe combinations are encoded differently in the distinct trial phases^63^. Furthermore, a recent study in monkeys suggested that the interplay between transient and stable coding guides the evolution of category based decisions during WM^34^. The advantage of a temporally stable representation during WM seems straightforward, also because such a low dimensional subspace could be read out over time by a set of fixed weights^33^. On the other hand, such a low dimensional representation would limit the information capacity for coding transient events. An intrinsic reservoir of continuous neuronal timescales therefore seems particularly well suited for parallelizing either computation, as implied within the present study. It will be important to further investigate the recruitment of neurons according to their intrinsic timescales when mnemonic information has to be integrated with flexible task demands.

Our initial quantification of baseline autocorrelation adds to the accumulating evidence for a hierarchy of intrinsic timescales in the monkey^36–38, 42&49^, mouse^39^ and human cortex^40, 41&64^. Prefrontal areas lPFC and FEF showed longer intrinsic timescales than LIP, which coincides with their position in the cortical hierarchy^65^. In the temporal domain, longer processing timescales directly relate to the integration or maintenance of information over time^39^, which is useful for higher cognitive functions such as decision making or WM^39, 46^. Indeed, prefrontal areas and specifically lPFC, which showed the longest average intrinsic timescale, has served as the major focus for WM research over the past decades^7^ and is generally considered crucial for WM function^66, 67^.

It is tempting to argue that neurons exhibiting longer intrinsic timescales form part of a continuous attractor network. Classical attractor models of WM persistent activity^18–21^ imply functional clustering i.e. increased connectivity between clusters of similarly tuned neurons, which has been observed and been linked to persistent activity in some tasks^68^. Local connection density per se might be a good predictor of intrinsic timescales and in turn persistent activity^8, 39&42^. In a recent study, Chaudhuri and colleagues constructed a large scale brain model aimed at investigating possible mechanisms leading to a hierarchy of intrinsic timescales similar to the one observed in^37, 42^. They found that both long range connections (‘inter-areal loops’) as well as local connection densities determined the relative size of intrinsic timescales across brain areas^37, 42^. In line with this, a recent study in mice found that areas with higher functional coupling showed larger processing timescales on the level of the population^39^. Together, those observations raise the possibility that an individual intrinsic timescale might be determined by the concentration of incoming and outgoing connections a given neuron receives i.e. whether it forms part of a local ‘network hub’^34^. Our results for FEF and LIP indeed suggest that neurons with longer intrinsic timescales might pertain to such a network hub. In lPFC we failed to observe a relationship between a neuron’s functional connectivity and intrinsic timescale, possibly indicating more flexible connectivity regimes^30, 69^. It is important to note that spike count correlations do not necessarily indicate anatomical connectivity and their source does not have to be local^54^. Finally, there exists a multitude of cell-intrinsic or network mechanisms underlying persistent activity accounts^10^. This plurality naturally applies to possible sources for the heterogeneity of observed intrinsic timescales.

Through analysing simultaneous electrophysiological recordings from three brain regions: lPFC, FEF and LIP, we found that the temporal tuning heterogeneity previously observed during WM is predicted by baseline firing stability of individual neurons. We showed that intrinsic timescales effectively shape the temporal dimensionality of the encoded information. In the prefrontal regions lPFC and FEF, neurons with longer intrinsic timescales encoded task information more strongly while storing it in a comparably stable, low dimensional format throughout the delay. In contrast, cells exhibiting short intrinsic timescales encoded relevant features in a more dynamic fashion, maintaining a high temporal coding dimensionality throughout the task, which in lPFC, could be robustly decoded even during the delay period. Additionally, in lPFC, the presented sample stimulus was more rapidly encoded by short timescale cells. Lastly, we presented some evidence for distinct mechanisms contributing to the emergence of intrinsic timescale heterogeneity. In summary, we suggest that WM constitutes the dynamic recruitment of neurons with different intrinsic timescales, possibly shaped by local connectivity regimes, to optimize functionally coexistent computations such as the encoding and storage of information.

## Methods

### Experimental paradigm and recordings

Physiological recordings have previously been reported in detail^51^. In brief, two adult rhesus monkeys (*Macaca mulatta*) were trained to perform a delayed change localization task (Fig. 1a). After a short fixation period (500ms) an array of colored squares was presented for 800ms (the sample period), followed by an 800-1000ms memory delay (delay period). Finally, a test array was presented to the animal. The test array was identical to the sample array except that one randomly chosen item changed color. To receive reward, monkeys had to make a saccade to the changed item. On each trial, the total number of items on the screen varied between two and five. There were six possible item locations and two possible color values per location (changing each session). Over several sessions, simultaneous recordings were taken from single neurons in the lateral prefrontal cortex (lPFC; 584 cells total); the frontal eye fields (FEF; 325 cells) and the lateral intraparietal cortex (LIP; 284 cells).

### Preprocessing

We excluded cells with fewer than 100 spikes per session from all analysis, hence reducing total cell counts per area to: 583 cells for lPFC, 323 cells for FEF and 281 cells for LIP. There were at least 20 trials per relevant stimulus condition. Unless otherwise noted, binary spike trains were convolved with a Gaussian kernel (s.d. = 20 ms) to produce firing rates in sp/s. For the crosstemporal discriminability analysis, firing rates were down-sampled from 1000 to 100 Hz, allowing for more efficient computing, while maintaining sufficiently high temporal resolution. All data analysis was implemented in Python using custom-written code.

### Statistical testing

Throughout this study, we used a non-parametric cluster-based permutation test^70^. In brief, the method compares some observed test statistic with a constructed null distribution while controlling for multiple comparisons across time. Null distributions were constructed in two ways: (1) By randomly permuting condition labels within cells 1000 times (PEV and across time discriminability analysis); (2) By randomly permuting intrinsic timescales across cells 1000 times for comparing median split intrinsic timescales (i.e. t-test or raw differences) and correlation coefficients. In general, the relevant test statistic or raw difference was computed for the observed data as well as for each of the 1000 permutations. This was done for each individual time point or pair of time points (across time discrimination analysis). For (1), points were classified by comparing the observed effect to the 95^th^ percentile of the null distribution. Contiguous points exceeding the 95^th^ percentile were deemed candidate clusters. For (2), cluster candidates were directly derived from the primary test statistic (contiguous analytical p-values < 0.05 from t-test or correlation analysis). To correct for multiple comparisons, we compared the maximum summed cluster test statistic of the observed data to a new null distribution of maximum summed clusters derived from our permutations. If the size of the observed cluster exceeded the 95^th^ percentile of the new null distribution it was deemed significant.

### Estimation of intrinsic timescales

#### Estimation of intrinsic timescales at the single cell level

Intrinsic timescales (τ) for single cells were estimated from the 500ms fixation period^37^. To calculate the temporal autocorrelation, spikes were counted in ten 50ms successive, independent time bins. This procedure created a trial by time-bin matrix. The temporal autocorrelation is the Pearson correlation across trials, between spike counts from each specific time-bin; i.e. we computed the correlation of each column of the matrix with each other. We fitted with an exponential decay function to the resulting autocorrelation time-course using non-linear least squares, as implemented by Levenberg-Marquardt algorithm within ScPy’s optimize_curve_fit function^37^ (Fig. 1b; Eq. 1).

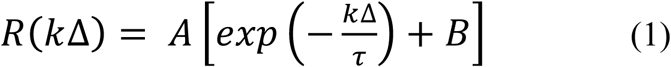

where **τ** is the intrinsic timescale, **A** is the amplitude and **B** is the offset parameter. The **k**Δ parameter refers to the relative time lag between time bins (50 – 450ms). Similar to previous studies^37, 49^, a fraction of cells showed relatively low autocorrelation values at short time lags, possibly due to refractory period or negative adaptation. To accommodate for that feature, fitting started at the first reduction in autocorrelation.

#### Exclusion criteria

Cells were not assigned an intrinsic timescale if they had: (1) less than 50 trials; (2) a fixation period firing rate lower than 1 spikes/s; (3) no spikes within any time bin across all trials or (4) first reduction in autocorrelation later than a time-lag of 150ms. Additionally, cells that were clearly not fit well by an exponential function, as determined by blinded visual inspection, were also not assigned a **τ** value. The remaining count of cells that were assigned a **τ** value and therefore were available for subsequent analysis was: 265 cells for lPFC, 168 cells for FEF and 136 cells for LIP.

### Information content of single cells

We based our estimation of single cell information content on the percentage of variance explained (ωPEV) statistic^51^. Specifically, we were interested in single cell selectivity to location and/or color information. First, we created a dummy coded design matrix where each column represented a stimulus condition, i.e. location one (1/6) and color one (1/2). For each trial, a one denoted that that specific condition was met whereas a zero denoted the opposite. We then performed a multiple linear regression with the above-defined conditions as independent factors and the firing rate as a dependent variable. Finally, we calculated ωPEV to quantify the modulation of individual firing rates accounted for by the combination of color and location of the presented stimulus items.

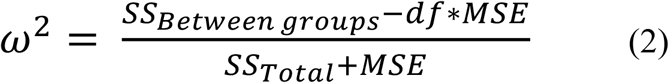

where **SS_Total_** denotes the total sum of squares i.e. total variance, **SS_Between Groups_** denotes the variance between groups, **df** are the degrees of freedom and **MSE** is the mean squared error of the model. Statistical significance of single cell PEVs was validated using cluster based permutation testing (see: *Statistical testing*) with randomly shuffled stimulus conditions.

### Cross temporal discriminability analysis

Cross temporal discriminability of stimulus information on the level of the neural population was assessed using the analysis described in the studies by Stokes et al. (2013)^27^ and Spaak et al. (2017)^28^. First, we randomly assigned each trial to one of two independent data splits. We then computed the mean firing rate over trials per neuron per independent split and per condition (locations * colors = 6 × 2 = 12) and took the pairwise differences between all conditions (66) for each neuron within each independent split. For each pairwise condition difference we then calculated the Pearson correlation for each time point across neurons between independent splits and averaged the resulting correlation coefficients using Fisher’s z-transformation to obtain a single time resolved discriminability measure. This procedure is analogous to a decoder being trained on split one and time point one (**t_1_**) and being tested on **t_1_**’ in split two. It is straightforward to extend this basic decoder to cross temporal decoding by computing the correlation between each time point (**t_n_**) in split one and all (same and other) time points **t_n+i_’** in split two. The result is a two dimensional matrix (time by time) representing stimulus discriminability at all time points (on diagonal) as well as across time points (off diagonals). Significance of discriminability was assessed by cluster-based permutation testing with randomly shuffled condition labels.

To decode location information only, we defined conditions with respect to the stimulus location, regardless of color (number of conditions = 6; pairwise differences = 15). To decode color information only, we focused on the difference between color conditions within each location (number of conditions = 12; within location differences = 6).

### Estimation of temporal coding dimensionality

#### Estimation of single cell temporal coding dimensionality

Principal component analysis (PCA) was used to estimate the temporal coding dimensionality of single cells. Commonly, PCA is used to explain the variance of population activity in neural state space in response to task features^53^. Here, we sought to apply the same idea to single neurons over time, thereby quantifying the temporal variability of task dependent firing. First, we constructed a data matrix **Y**: For each trial, we counted raw spikes falling into independent 50 ms time windows (spanning the 2000 ms trial). We then averaged binned spike counts by task condition (locations * colors = 6 × 2 = 12). Columns in **Y** correspond to 10 adjacent, independent time windows (spanning 500ms), (i.e. dimensions), and rows to the 12 task condition averages (i.e. samples). PCA is then performed on **Y** to quantify and arrange the variance along orthogonal axes i.e. principal components in the ‘space’ spanned by the ten independent time bins. The eigenvalues associated with each principal component give us the means to quantify the effective dimensionality (N*_eff_*)^71^ of our temporal state space.

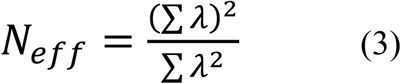

where **λ** represent the eigenvalues. The N_*eff*_ measure penalizes small eigenvalues which could arise due to noise. A high N_*eff*_ (the maximum equals ten in our example) suggests a high independence across time, i.e. a high temporal coding dimensionality. In other words, a cell with a high temporal dimensionality has either an unstable temporal condition selectivity (i.e., noise) or switches its condition selectivity over time.

### Temporal discriminability analysis

To decode task information in single cells by leveraging variability over time, we applied a temporal discriminability analysis described in the following. For each neuron, we randomly assigned each trial to one of two independent data splits. We then computed the mean firing rate over trials per independent split and per condition (locations * colors = 6 × 2 = 12) and took the pairwise differences between all conditions (66) within each independent split. For each pairwise difference, we computed the Pearson correlation between time-points from a 200ms window in split A of the data with the same time points in split B of the data. This was done for all pairwise condition differences and the resulting correlation coefficients were averaged using using Fisher’s z-transformation. By sliding the 200ms window along the trials (1 ms increments) and repeating the analysis we obtained a temporal discriminability score, which captures the information present in the temporal structure of the signal. Temporal variability that is purely driven by noise will result in a low temporal variability score, as will condition specific but stable firing within the 200ms. Conversely, dynamic but specific firing will result in a comparably high temporal discriminability score.

### Computing spike count correlations

Spike count correlations (rSC) (also known as noise correlations) (Cohen and Kohn, 2011) were calculated from the 500ms fixation period. We derived rSC values by comparing single cell responses recorded form the same electrode on the same session. This criterion limited our analysis to electrodes on which more than one cell was recorded. Furthermore, not all cells recorded by an electrode had the same number of trials, hence, we were limited to trials common to all cells recorded on one electrode. Here, we set the minimum inclusion criterion to 20 trials. Lastly, we only included cells that had an average firing rate greater than 2 sp/s during the fixation period. Together with our sub-selection of cells that were assigned a τ value we were left with: 126 cells for lPFC; 74 cells for FEF and 55 cells for LIP.

For each trial, we counted all spikes falling into the 500ms fixation period window. We then normalized spike rates on each trial for both mean firing and slow drifts in neural excitation by computing a z-score for each neuron’s firing rate on each trial using a sliding window of 10 trials before and after the current trial^72^.

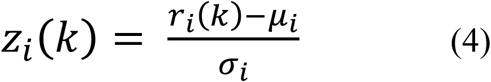

where **r** is the firing rate at trial **k** and **μ** and **σ** are the mean and standard deviation of the **i**^th^ neuron’s firing rate estimated from the 21 trials centered on **k**. The Pearson correlation of a neuron’s z-scores with those of any other neuron give the rSC. Ultimately, our goal was to correlate rSC with τ. Therefore, in addition to its τ value, each cell was assigned the Fisher-transformed average rSC of all its pairings on the same contact. If only two cells were recorded on one electrode, we reassigned both cells the mean of their respective log transformed τ values. Furthermore, to evaluate whether a possible correlation between τ and rSC was confounded by firing rates^55^, we also took the mean of the log transformed baseline firing rates making up each averaged rSC value.

## Acknowledgements

This research was funded by Biotechnology & Biological Sciences Research Council (BB/M010732/1) to MGS, ONR (N00014-14-1-0681) and NIMH (R00MH092715) to TJB, and NIMH (R37MH087027), The MIT Picower Institute Innovation Fund and ONR MURI grant (N00014-16-1-2832) to EKM, and was supported by the NIHR Oxford Health Biomedical Research Centre. The Wellcome Centre for Integrative Neuroimaging is supported by core funding from the Wellcome Trust (203139/Z/16/Z).

## Supplementary information

**Supplementary Figure 1:**
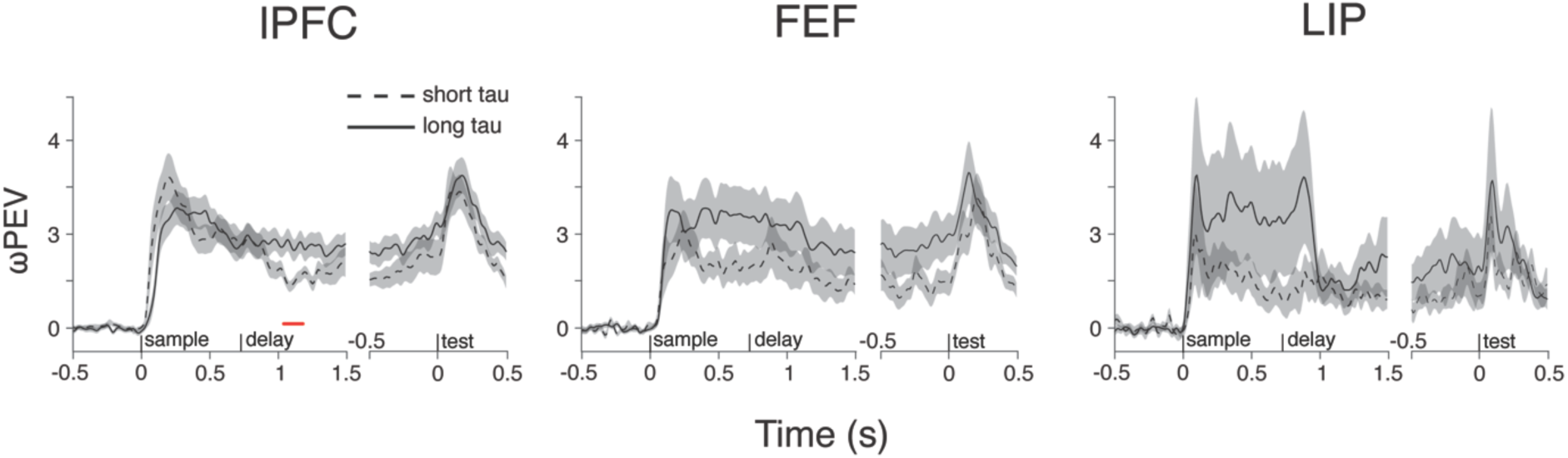
Item information and intrinsic timescales for cells showing a significant ωPEV. Same conventions as in Fig. 2 hold true. Red line indicates a significant difference (p=0.04).

**Supplementary Figure 2:**
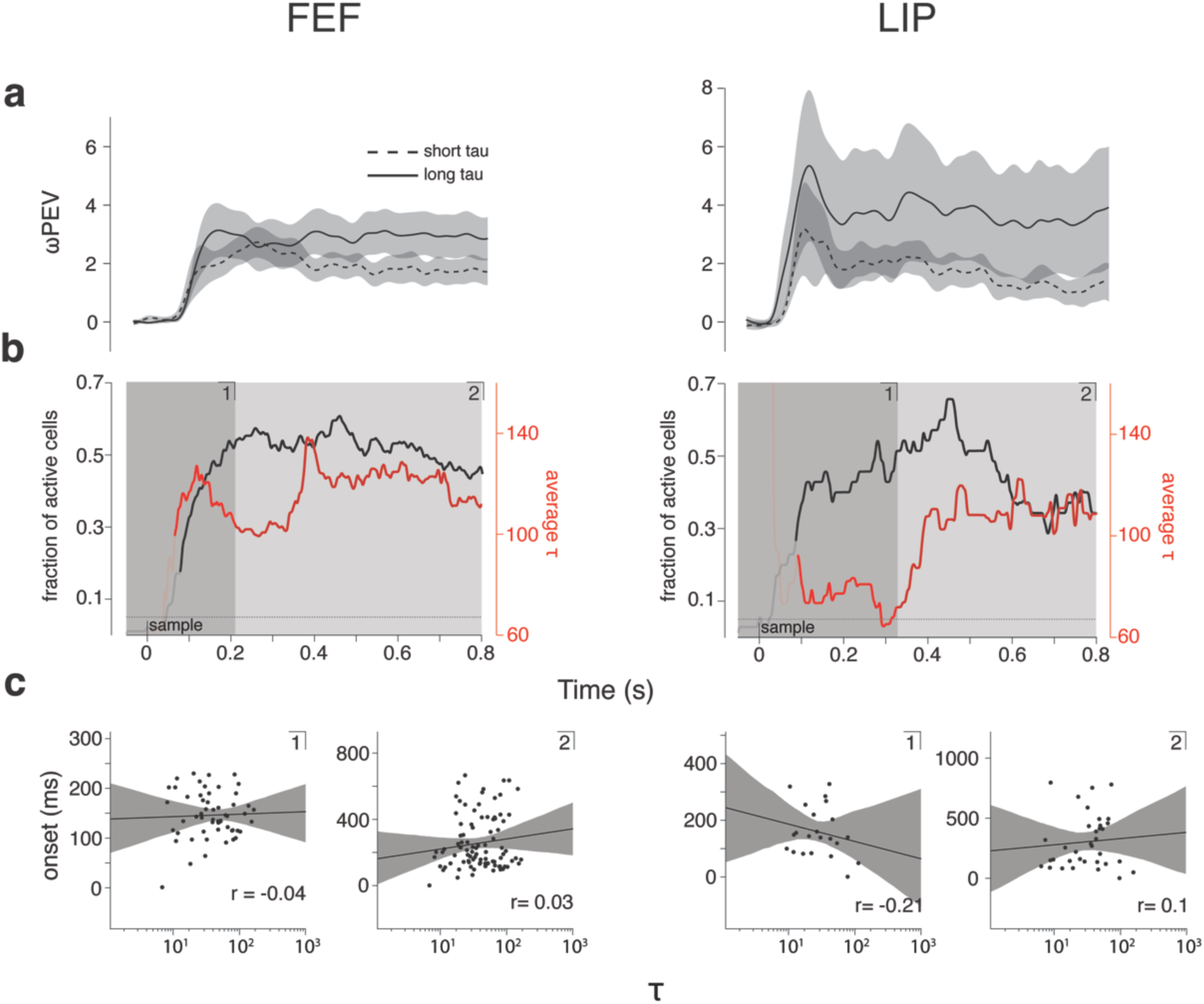
Evaluation of sample period onset times in relation to intrinsic timescales for FEF and LIP. This figure shows the same plots as Fig. 3, for FEF (left column) and LIP (right column). FEF: Time period **1** (Spearman correlation: r=-0.04, p=0.76, n=56); Time period **2** (Spearman correlation: r=0.04, p=0.71, n=96). LIP: Time Period **1**: (Spearman correlation: r=-0.21, p=0.34, n=22); Time Period **2**: (Spearman correlation: r=0.1, p=0.57, n=35).

**Supplementary Figure 3:**
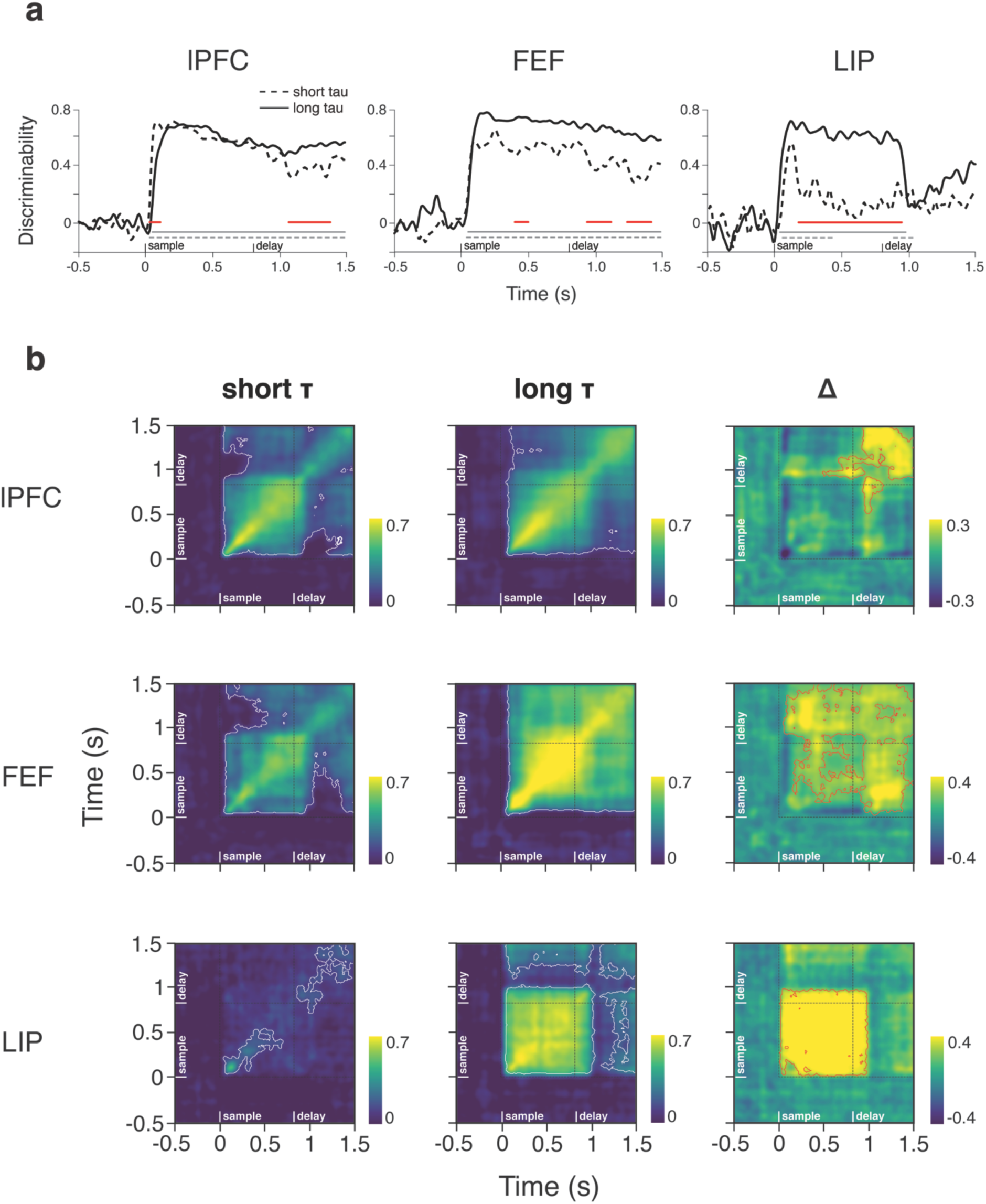
Within-time and cross-temporal decoding for location information. Same as Fig. 4 but for location information only (see Methods). White contours: p<0.001; Red contours: p<0.01.

**Supplementary Figure 4:**
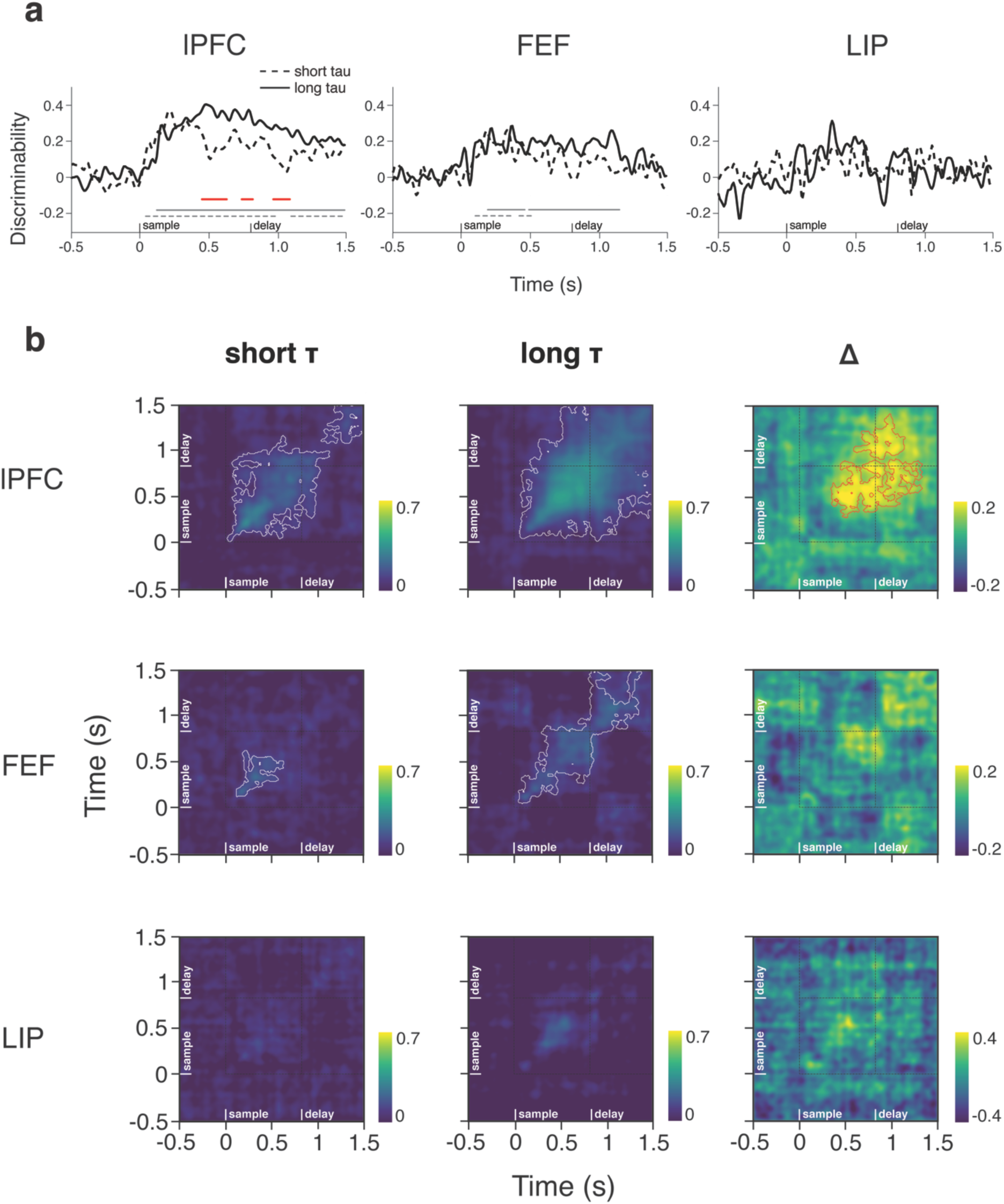
Within-time and cross-temporal decoding for color information. Same as Fig. 4 but for color information only (see Methods). White contours: p<0.001; Red contours: p<0.01.

**Supplementary Figure 5:**
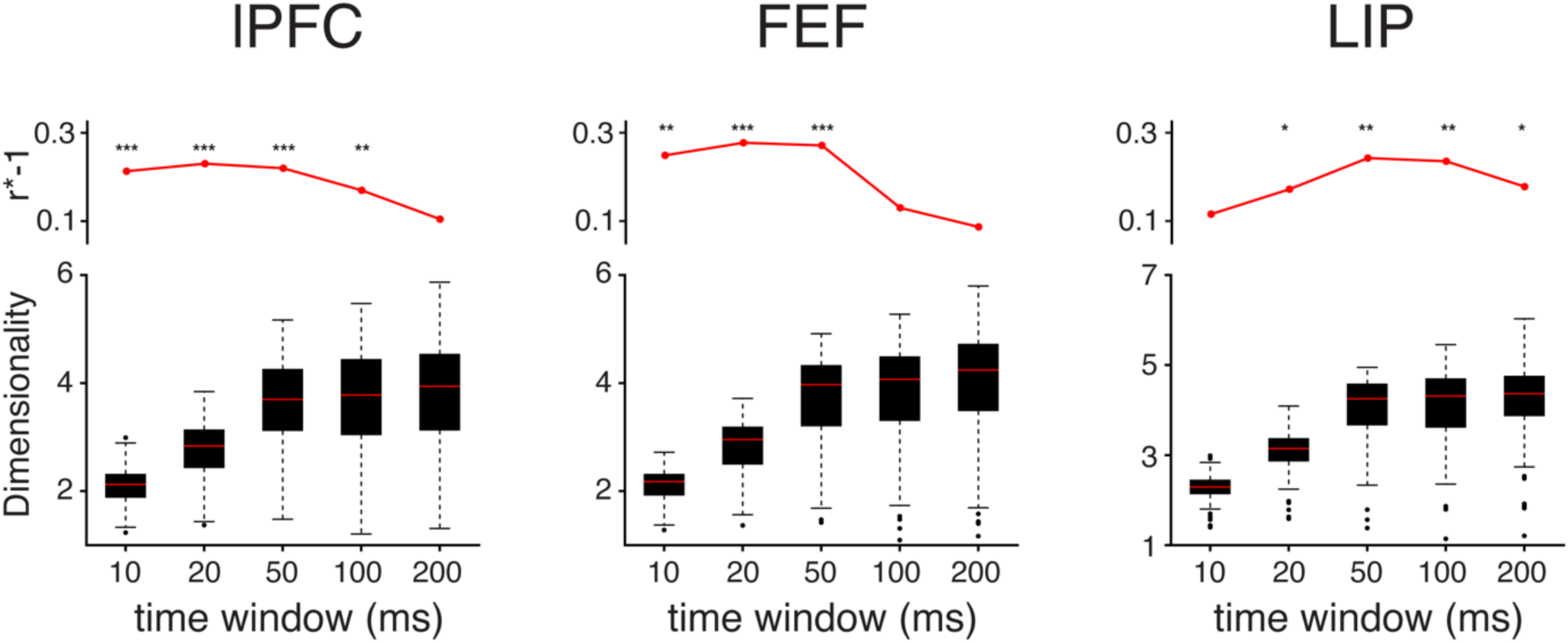
Effective temporal dimensionality estimated over different time windows and relationship with intrinsic timescales. Lower panels: Box-Whisker plots show N_*eff*_ estimated over differing timewindows (10ms*10, 20ms*10, 50ms*10, 100ms*10 and 200ms*10). Red lines denote median N_*eff*_ for each timewindow over all cells, black boxes mark the upper and lower quartiles. Dots represent outliers. Upper panels: Each dot represents the Spearman correlation coefficient of N_*eff*_ and τ vlues, for each of the time window N_*eff*_ is estimated from. * represents p<0.05; ** represents p<0.01; *** represents p<0.001. lPFC (n=264); FEF (n=167); LIP (n=134).

**Supplementary Figure 6:**
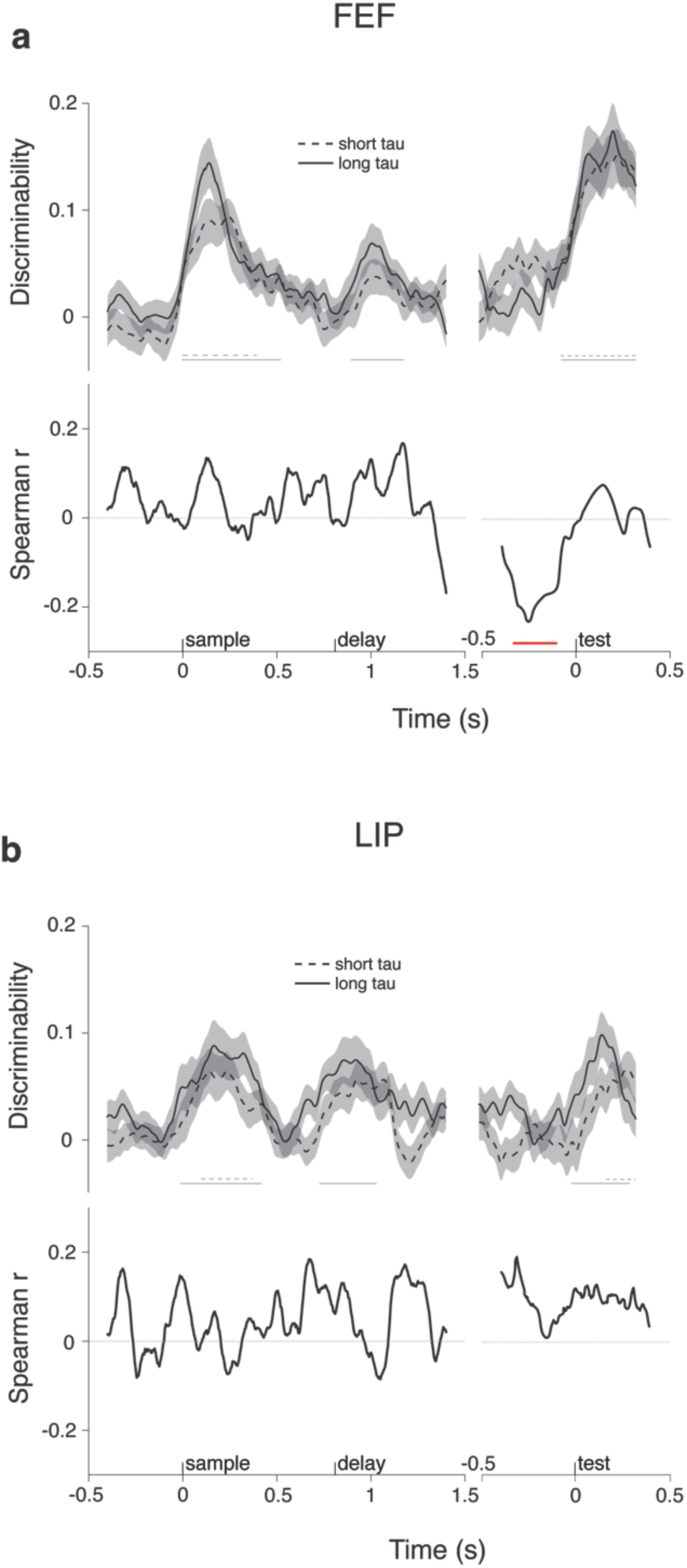
Temporal discriminability in FEF and LIP. Same conventions as in Fig. 6. **a)** FEF; **b)** LIP. Red line indicated a significant difference (cluster based permutation test; p<0.05)

**Supplementary Figure 7:**
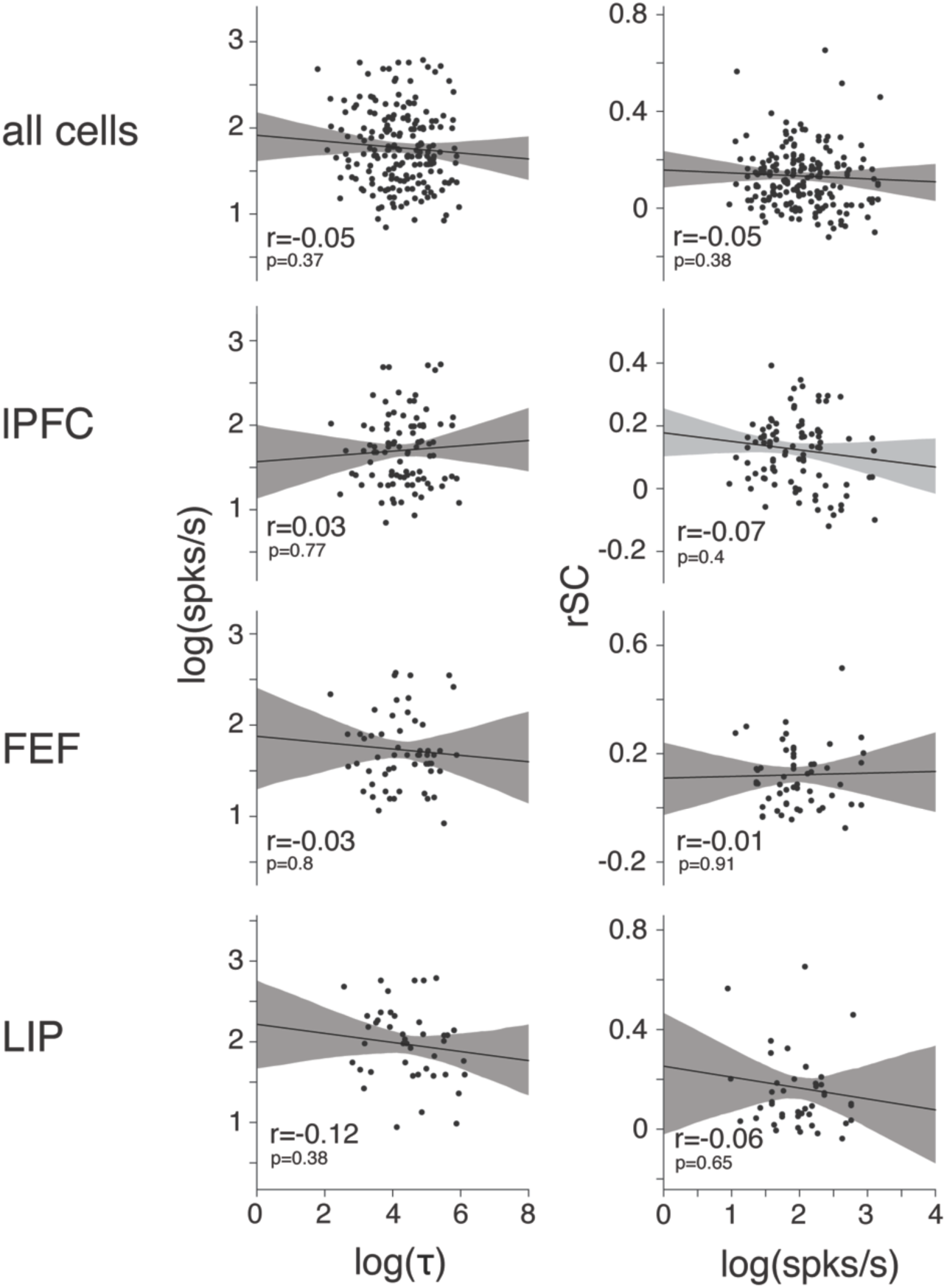
Spike count correlations and intrinsic timescales. The first column shows scatter plots for the log transformed fixation period firing rate in spks/s on the y-axis, versus the log transformed τ values on the x-axis. The r value within panels depicts the Spearman correlation coefficient of the two variables. Black solid lines show linear regression lines. Shaded region depicts 95% confidence interval. All cells: r=-0.07, p=0.29, n=255; lPFC: r=0.05, p=0.54, n=126; FEF: r=0.07, p=0.58, n=74; LIP: r=-0.12, p=0.4, n=55. The third column shows scatter plots for the rSC on the y-axis, versus the log transformed fixation period firing rate (spks/s) on the x-axis. All cells: r=-0.05, p=0.45, n=255; lPFC: r=-0.12, p=0.19, n=126; FEF: r=0.02, p=0.84, n=74; LIP: r=-0.12, p=0.38, n=55. The r value within panels depicts the Spearman correlation coefficient and associated p-value, between the two variables

## References

1. Rigotti, M., Barak, O., Warden, M. R., Wang, X. J., Daw, N. D., Miller, E. K., Fusi, S. The importance of mixed selectivity in complex cognitive tasks. Nature 497, 585–90 (2013).

2. Raposo, D., Kaufman, M. T., Churchland, A. K. A Category-free neural P population supports evolving demands during decision-making. Nat. Neurosci. 17, 1784–92 (2014).

3. Funahashi, S., Bruce, C. J., Goldman-Rakic, P. S. Mnemonic coding of visual space in the monkey’s dorsolateral prefrontal cortex. J. Neurophysiol. 61, 331–49 (1989).

4. Miller, E. K., Erickson, C. A., Desimone, R. Neural mechanisms of visual working memory in prefrontal cortex of the macaque. J. Neurosci. 16, 5154–67 (1996).

5. Fuster, J. M. Network memory. Trends Neurosci. 20, 451–459 (1997).

6. Fuster, J. M., and Alexander, G. E. Neuron activity related to short-term memory. Science 173, 652–54, (1971).

7. Goldman-Rakic, P. S. Cellular basis of working memory. Neuron 14, 477–85 (1995).

8. Barak, O., and Tsodyks, M. Working models of working memory. Curr. Opin. Neurobiol. 25, 20–24 (2014).

9. Stokes, M. G. ‘Activity-silent’ working memory in prefrontal cortex: A dynamic coding framework. Trends Cogn. Sci. 19, 394–405 (2015).

10. Zylberberg, J., and Strowbridge, B. W. Mechanisms of persistent activity in Cortical circuits: Possible neural substrates for working memory. Annu. Rev. Neurosci. 40, 603–27 (2017).

11. Funahashi, S., Bruce, C. J., Goldman-Rakic, P. S. Mnemonic coding of visual space in the monkey’s dorsolateral prefrontal cortex. J. Neurophysiol. 61, 331–49 (1989).

12. Chafee, M. V. and Goldman-Rakic, P. S. Matching patterns of activity in primate prefrontal area and parietal area neurons during a spatial working memory task. J. of Neurophysiol. 79, 2919–40 (1998).

13. Rainer, G., Asaad, W. F., Miller, E. K. Memory fields of neurons in the primate prefrontal cortex. PNAS 95, 15008–13 (1998).

14. Romo, R., Brody, C. D., Hernández, A., Lemus, L. Neuronal correlates of parametric working memory in the prefrontal cortex. Nature 399, 470–73 (1999).

15. Loewenstein, Y., and Sompolinsky, H. Temporal integration by calcium dynamics in a model neuron. Nat. Neurosci. 6, 961–67 (2003).

16. Fransén, E., Tahvildari, B., Egorov, A. V., Hasselmo, M. E., Alonso, A. A. Mechanism of graded persistent cellular activity of entorhinal cortex layer V neurons. Neuron 49, 735–46 (2006).

17. Amit, D. J., and Brunel N. Model of global spontaneous activity and local structured activity during delay periods in the cerebral cortex. Cerebr. Cortex 7, 237–52 (1997).

18. Compte, A., Brunel, N., Goldman-Rakic, P. S., Wang, X. J. Synaptic mechanisms and network dynamics underlying spatial working memory in a cortical network model. Cerebr. Cortex 10, 910–23 (2000).

19. Wang, X. J. Synaptic Reverberation underlying mnemonic persistent activity. Trends Neurosci. 24, 455–63 (2001).

20. Machens, C. K., Romo, R., Brody, C. D. Flexible control of mutual inhibition: A neural model of two-interval discrimination. Science 307, 1121–24 (2005).

21. Wong, K. F., and Wang, X. J. A Recurrent network mechanism of time integration in perceptual decisions. J. of Neurosci. 26, 1314–28 (2006).

22. Brody, C. D., Hernández, A., Zainos, A., Romo, R. Timing and neural encoding of somatosensory parametric working memory in macaque prefrontal cortex. Cerebr. Cortex 13, 1196–1207 (2013).

23. Shafi, M., Zhou, Y., Quintana, J., Chow, C., Fuster, J., Bodner, M. Variability in neuronal activity in primate cortex during working memory tasks. Neurosci. 146, 1082–1108 (2007).

24. Enel, P. Procyk, E., Quilodran, R., Dominey, P. F. Reservoir computing properties of neural dynamics in prefrontal cortex. PLOS Comput. Biol. 12, e1004967 (2016).

25. Lundqvist, M., Rose, J., Herman, P., Brincat, S. L., Buschman, T. J., Miller, E. K. Gamma and beta bursts underlie working memory. Neuron 90, 152–64 (2016).

26. Meyers, E. M., Freedman, D. J., Kreiman, G., Miller, E. K., Poggio, T. Dynamic population coding of category information in inferior temporal and prefrontal cortex. J. of Neurophysiol. 100, 1407–19 (2008).

27. Stokes, M. G., Kusunoki, M., Sigala, N., Nili, H., Gaffan, D., Duncan, J. Dynamic coding for cognitive control in prefrontal cortex. Neuron 78, 364–75 (2013).

28. Spaak, E., Watanabe, E., Funahashi, S., Stokes, M. G. Stable and dynamic coding for working memory in primate prefrontal cortex. J. of Neurosci. 37, 6503–16 (2017).

29. Maass, W., Natschläger, T., Markram, H. Real-time computing without stable states: A new framework for neural computation based on perturbations. Neural Comput. 14, 2531–60 (2002).

30. Mongillo, G., Barak, O., Tsodyks, M. Synaptic theory of working memory. Science 319, 1543–46 (2008).

31. Druckmann, S., and Chklovskii, D. B. Neuronal circuits underlying persistent representations despite time varying activity. Curr. Biol. 22, 2095–2103 (2012).

32. Barak, O., Sussillo, D., Romo, R., Tsodyks, M., Abbott, L. F. From fixed points to chaos: Three models of delayed discrimination. Prog. Neurobiol. 103, 214–22 (2013).

33. Murray, J. D., Bernacchia, A., Roy, N. A., Constantinidis, C., Romo, R., Wang, X. J. Stable population coding for working memory coexists with heterogeneous neural dynamics in prefrontal cortex. PNAS 114, 394–99 (2017).

34. Chaisangmongkon, W., Swaminathan, S. K., Freedman, D. J., Wang, X. J. Computing by Robust Transience: How the fronto-parietal network performs sequential, category-based decisions. Neuron 93, 1504–1517 (2017).

35. Hubel, D. H., and Wiesel, T. N. Receptive fields, binocular interaction and functional architecture in the cat’s visual cortex. J. of Physiol. 160, 106–154 (1962).

36. Chen, J., Hasson, U., Honey, C. J. Neuron 88, 244–46 (2014).

37. Murray, J. D., Bernacchia, A., Freedman, D. J., Romo, R., Wallis, J. D., Cai, X., Padoa-Schioppa, C., Wang, X. J. A Hierarchy of intrinsic timescales across primate cortex. Nat. Neurosci. 17, 1661–63 (2014).

38. Ogawa, T., and Komatsu, H. Differential temporal storage capacity in the baseline activity of neurons in macaque frontal eye field and Area V4. J. of Neurophysiol. 103, 2433–45 (2010).

39. Runyan, C. A., Piasini, E., Panzeri, S., Harvey, C. D. Distinct timescales of population coding across cortex. Nature 548, 92–96 (2017).

40. Hasson, U., Yang, E., Vallines, I., Heeger, D. J., Rubin, N. A hierarchy of temporal receptive windows in human cortex. J. Neurosci. 28, 2539–50 (2008).

41. Honey, C. J., Thesen, T., Donner, T. H., Silbert, L. J., Carlson, C. E., Devinsky, O., Doyle, W. K., Rubin, N., Heeger, D. J., Hasson, U. Slow cortical dynamics and the accumulation of information over long timescales. Neuron 76, 423–34 (2012).

42. Chaudhuri, R., Knoblauch, K., Gariel, M. A., Kennedy, H., Wang, X. J. A large-scale circuit mechanism for hierarchical dynamical processing in the primate cortex. Neuron 88, 419–31 (2015).

43. Goldman, M. S., Compte, A., Wang, X. J. Encyclopedia of Neuroscience (Ed. Squire L.R.). 165–178 (Academic Press, Oxford, 2008).

44. Buračas, G. T., Zador, A. M., DeWeese, M. R., Albright, T. D. Efficient discrimination of temporal patterns by motion-sensitive neurons in primate visual cortex. Neuron 20, 959–69 (1998).

45. Salinas, E., Hernández, A., Zainos, A., Romo, R. Periodicity and firing rate as candidate neural codes for the frequency of vibrotactile stimuli. J. Neurosci. 20, 5503–15 (2000).

46. Wang, X. J. The Prefrontal Cortex as a quintessential "cognitive-type" neural circuit. In Principles of Frontal Lobe Function, edited by Stuss D. T. and Knight, R. T. (Oxford University Press, Oxford, 2013).

47. Scott, B. B, Constantinople, C. M., Akrami, A., Hanks, T. D., Brody, C. D., Tank, D. W. Fronto-parietal cortical circuits encode accumulated evidence with a diversity of timescales. Neuron 95, 385–398 (2017).

48. Bernacchia, A., Seo, H., Lee, D., Wang, X. J. A Reservoir of time constants for memory traces in cortical neurons. Nat. Neurosci. 14, 366–72 (2013).

49. Cavanagh, S. E., Wallis, J. D., Kennerley, S. W., Hunt, L. T. Autocorrelation structure at rest predicts value correlates of single neurons during reward-guided choice. eLife 5, e18937 (2016).

50. Nishida, S., Tanaka, T., Shibata, T., Ikeda, K., Aso, T., Ogawa, T. Discharge-rate persistence of baseline activity during fixation reflects maintenance of memory-period activity in the macaque posterior parietal cortex. Cerebr. Cortex 24, 1671–85 (2014).

51. Buschman, T. J., Siegel, M., Roy, J. E., Miller, E. K. Neural substrates of cognitive capacity limitations. PNAS 10, 11252–55 (2011).

52. Sigala, N., Kusunoki, M., Nimmo-Smith, I., Gaffan, D., Duncan, J. Hierarchical Coding for Sequential Task Events in the Monkey Prefrontal Cortex. PNAS 105, 11969–74 (2008).

53. Cunningham, J. P., and Yu, B. M. Dimensionality Reduction for Large-Scale Neural Recordings. Nat. Neurosci. 17, 1500–1509 (2014).

54. Cohen, M. R. and Kohn A. Measuring and interpreting neuronal correlations. Nat. Neurosci. 14, 811–19 (2011).

55. De la Rocha, J., Doiron, B., Shea-Brown, E., Josic, K., Reyes, A. Correlation between neural spike trains increases with firing rate. Nature 448, 802–6 (2007).

56. Cromer, J. A., Roy, J. E., Miller, E. K. Representation of multiple, independent categories in the primate prefrontal cortex. Neuron 66, 796–807 (2010).

57. Koyluoglu, O. O., Pertzov, Y., Manohar, S., Husain, M., Fiete, I. R. Fundamental bound on the persistence and capacity of short-term memory stored as graded persistent activity. eLife, doi:10.7554 (2017).

58. Markowitz, D. A., Curtis, C. E., Pesaran, B. Multiple component networks support working memory in prefrontal cortex. PNAS 112, 11084–89 (2015).

59. Pinto, L., and Dan, Y. Cell-type-specific activity in prefrontal cortex during goal-directed behavior. Neuron 87, 437–50 (2015).

60. Mendoza-Halliday, D., and Martinez-Trujillo, J. C. Neuronal population coding of perceived and memorized visual features in the lateral prefrontal cortex. Nat. Commun. 8, doi:10.1038 (2017).

61. Stokes, M. J. Neurosci. 31, 1167–69 (2011).

62. Duncan, J., and Miller, E. K. Cognitive focus through adaptive neural coding in the primate prefrontal cortex. Principles of Frontal Lobe Function, 278–91 (Oxford University Press, Oxford, 2002).

63. Blackman, R. K., Crowe, D. A., DeNicola, A. L., Sakellaridi, S., MacDonald, A. W., Chafee, M. V. Monkey Prefrontal neurons reflect logical operations for cognitive control in a variant of the AX continuous performance task (AX-CPT). J. Neurosci. 36, 4067–79 (2016).

64. Gauthier, B., Eger, E., Hesselmann, G., Giraud, A. L., Kleinschmidt, A. Temporal tuning properties along the human ventral visual stream. J. Neurosci. 32, 14433–41 (2012).

65. Felleman, D. J., and Van Essen, D. C. Distributed hierarchical processing in the primate cerebral cortex. Cerebr. Cortex 1, 1–47 (1991).

66. Butters, N., and Pandya, D. Retention of delayed-alternation: Effect of selective lesions of sulcus principalis. Science 165, 1271–73 (1969).

67. Stamm, J. S. Electrical stimulation of monkeys’ prefrontal cortex during delayed-response performance. J. Comp. Physiol. Psychol. 67, 535–46 (1969).

68. Masse, N. Y., Hodnefield, J. M., Freedman, D. J. Mnemonic Encoding and Cortical Organization in Parietal and Prefrontal Cortices. J. Neurosci. 37, 6098–6112 (2017).

69. Szatmáry, B., and Izhikevich, E. M. Spike-timing theory of working memory. PLOS Comput. Biol. 6, e1000879 (2010).

70. Maris, E., and Oostenveld, R. Nonparametric statistical testing of EEG-and MEG-data. J. Neurosci. Methods 164. 177–90 (2007).

71. Abbott, L. F., Rajan, K., Sompolinsky, H. Interactions between intrinsic and stimulus-evoked activity in recurrent neural networks. In: Neuronal variability and its functional significance (Oxford University Press, Oxford, 2010)

72. Cohen, M. R. and Newsome, W. T. Context-dependent changes in functional circuitry in visual area MT. Neuron 60, 162–73 (2008).

73. Rose S. N., La Rocque, J. J., Riggall, A. C., Gosseries, O., Starrett, M. J., Meyering, E. E., Postle, B. R. Reactivation of latent working memories with transcranial magnetic stimulation. Science 354, 1136–1139 (2016).

